# Tamoxifen Targets Wisp2 to Impair Subcutaneous Adipocyte Progenitor Self-Renewal and Adipogenic Differentiation

**DOI:** 10.64898/2025.12.21.695772

**Authors:** Nisha S. Thomas, Barbara Mensah Sankofi, Haoning Howard Cen, Stevi J. Murguia, Michael C. Rudolph, Elizabeth A. Wellberg

## Abstract

Breast cancer endocrine therapy, which systemically disrupts estrogen receptor signaling, increases type 2 diabetes (T2D) risk in some women. Sustained treatment with low-dose tamoxifen depletes subcutaneous adipocyte progenitors and promotes glucose intolerance and hepatic lipid deposition in obese female mice. Hyperplastic adipose tissue expansion, especially in subcutaneous depots, preserves metabolic health during a chronic positive energy balance by facilitating nutrient storage and attenuating inflammation. Adipocyte progenitors are renewed in part through Wnt signaling pathway activation, which is altered in women with obesity or T2D. Estrogen receptors are expressed in several adipose cell types, but the distinct actions of tamoxifen in adipocyte progenitors and the mechanisms that explain their depletion during endocrine therapy are not defined. The direct impact of tamoxifen was evaluated in subcutaneous adipose stromal cells from humans and adult mice. Self-renewal, proliferation, and differentiation were measured, and analyses of gene expression and progenitor or preadipocyte populations were performed. Mechanistic insight was gained from primary adipose stromal cells of obese female mice, in which the Wnt1 inducible signaling pathway protein 2 (Wisp2) was lost following endocrine therapy. Wisp2 gain and loss of function studies were carried out in adipose stromal cells to define the link between estrogen signaling and adipocyte progenitor maintenance. We found that tamoxifen treatment disrupts the protection of adipocyte progenitors by estrogen, mediated through suppression of Wisp2. These studies reveal potential metabolic effects of tamoxifen therapy that precede and could drive T2D development in breast cancer survivors.

## INTRODUCTION

Estrogen Receptor (ER) positive breast cancer, which accounts for 70% of all breast cancer cases, has seen significant improvements in patient survival due to early detection and the advancements in therapeutic approaches, particularly endocrine therapy. However, these lifesaving therapies are associated with elevated risk of developing early onset or subclinical metabolic complications in some patients (1, 2). Endocrine therapies include selective ER modulators like tamoxifen or aromatase inhibitors, and function by either blocking ovarian estrogen production or antagonizing estrogen receptor alpha (ERα) activity in breast cancer cells. These systemic treatments also block ER signaling in metabolic tissues including adipose, liver, and skeletal muscles, with the potential to disrupt whole body metabolism (3). Clinical studies report an increased risk of some metabolic disruptions in breast cancer patients following endocrine therapy, including insulin resistance, hepatic steatosis, and weight gain (4–16). Preclinical data, including our previous study, suggest that adipose tissue is a primary mediator of metabolic dysfunction during endocrine therapy (17–20). Estrogen and ER influence regional adipose distribution and function in depot- and sex-dependent manners. Within adipose stromal fractions are adipocyte precursor cells (APCs), which can be classified as progenitors or committed preadipocytes based on cell surface markers such as Sca1 and CD24 (21). Estrogen signaling enhances hyperplastic expansion of adipose tissue, especially during positive energy balance (22–25). Renewal and maintenance of APCs through proliferation is a prerequisite for healthy adipose tissue expansion to temper adipocyte hypertrophy that can promote ectopic fat deposition and insulin resistance. We previously found that tamoxifen and estrogen deprivation treatments, which simulate breast cancer endocrine therapy, prevented adipocyte progenitor expansion during a positive energy balance, and promoted adipocyte hypertrophy, glucose intolerance, and hepatic steatosis in obese female mice (26). Here, we show that estradiol (E_2_) and tamoxifen directly influence adipocyte progenitor expansion and differentiation. We identify the Wnt-1 inducible signaling pathway protein 2 (Wisp2) as a key mediator of the response of adipocyte precursor cells to estrogen and anti-estrogen signaling. Wisp2 has documented roles in the proliferation and maintenance of adipocyte progenitors. *In vivo*, Wisp2 loss from adipose stromal cells leads to hypertrophic adipose tissue growth during obesity development, accompanied by liver fat deposition and insulin resistance (27). Conversely, overexpression of Wisp2 in adipocyte precursors facilitates their proliferation and reduces unhealthy adipose tissue expansion (28). We demonstrate that, within the heterogeneous populations of subcutaneous adipose stromal cells, Wisp2 integrates the opposing effects of estrogen and tamoxifen on progenitor pools, serving as a potential mechanism through which endocrine therapy interferes with hyperplastic adipose tissue growth during weight gain.

## MATERIALS & METHODS

### Adipose Stromal Cell Isolation and Culture

Mouse immortalized subcutaneous adipocyte precursors (APCs; Kerafast #EVC005) from adult C57Bl/6 male mice, were cultured in 1x DME/F-12 medium supplemented with 10 % FBS, 1% Pen/Strep, and 1% sodium pyruvate. The cells were passaged using 0.25% Trypsin/EDTA every 3 days. All cell culture reagents were obtained from Thermo-Fisher.

Wisp2 knockdown (W2KO) cells were generated using mouse WISP-2 shRNA lentiviral particles (Santa Cruz Biotechnology) with 5 μg/ml polybrene. Control cells were transduced with GFP shRNA (Santa Cruz Biotechnology). Transduced cells were selected using 1 μg/mL puromycin. All cell lines were maintained under sterile conditions at 37°C and 5% CO_2._ All hormone treatments were performed in phenol red-free 1x DMEM (Gibco) containing 0.5% charcoal-stripped FBS and 1% Pen/Strep.

#### Primary mouse adipose stromal cell isolation

Inguinal subcutaneous adipose tissues from adult wildtype C57BL/6J female mice were digested in warm 1x HBSS containing 3% FBS, 40 ug/mL Pen/Strep, 40 ug/mL gentamycin, 500 ng/mL Amphotericin B, and 1 mg/mL collagenase I (Worthington) and incubated in a shaker (100 rpm) at 37⁰C for 1 hour. Tubes were inverted after 30 minutes to ensure proper digestion. The reaction was stopped with 10 mL cold 1x HBSS with 10% FBS. Cell suspensions were passed through 40 μm strainer and centrifuged at 1500 rpm for 5 minutes to liberate the stromal fraction. Pellets were resuspended in 500 µL red blood cell lysis buffer. 10 mL 1x HBSS was added and cells were centrifuged at 1500 rpm for 5 minutes. Resulting adipose stromal cells (ASCs) were resuspended in 1x DMEM with 10% FBS and 1% Pen/Strep.

To generate Wisp2 overexpressing cells (W2OE), primary ASCs were isolated from subcutaneous white adipose tissue of *Ccn5*-ROSA26 transgenic mice (Supplemental Fig 1) and transduced with adenoviral Cre recombinase (Ad-CMV-Cre; Vector Biolaboratories) or control adenovirus (Ad-GFP; Vector Biolaboratories) at MOI of 200. ASCs were incubated with virus-containing media for 6 hours, followed by a wash with 1x PBS. The cells were suspended in standard growth medium.

#### Serial passaging of primary mouse ASCs

ASCs from six C57BL/6 females were isolated as described above. 0.5 million cells were plated in 1x DMEM containing 5% charcoal-stripped FBS with EtOH vehicle (Veh), E_2_ or E_2_+TAM treatments. At 80% confluence, cells were trypsinized, counted, and 0.5 million viable cells from each group were replated with treatments. 0.1 million cells from each treatment per passage were plated for adipogenesis assays. Cell differentiation was assessed using oil red O staining.

#### Primary human precursor culture and maintenance

Primary human subcutaneous preadipocytes (PCS-210-01, ATCC) were cultured in fibroblast basal medium (ATCC PCS-201-030) with fibroblast low serum growth supplement (ATCC PCS-201-041). Cells were passaged using 0.05% trypsin-EDTA. For adipogenic differentiation, once cells reached confluence, adipocyte differentiation medium (ATCC PCS-500-050) was added for 4 days. After 4 days, cells were further cultured in adipocyte basal medium supplemented with adipocyte maturation agents (ADM, ATCC PCS-500-050). On day 15, adipocyte phenotype and lipid accumulation were evaluated by oil Red O staining All experimental conditions were conducted in triplicate, with differentiation experiments independently repeated a minimum of three times.

All cells were treated with one or more of the following compounds purchased from Sigma: 10nM 17β-estradiol (E_2_); 100pM 4-OH tamoxifen (TAM); 0.001% EtOH (Veh). Hormone treatments were performed in phenol red-free 1x DMEM supplemented with 0.5% charcoal stripped FBS and 1% Pen/Strep.

### Adipocyte Differentiation

Differentiation was initiated in confluent primary ASCs or immortalized APCs (day 0) with complete media (1x DMEM/F-12 for immortalized APCs and 1x DMEM for primary APCs) containing 0.5 mM 3-Isobutyl-1-methylxantithine (IBMX), 1 µM dexamethasone, 10 µg/mL insulin and 1 µM rosiglitazone for 48 hours. Cells were then switched to maintenance media containing 10 µg/ml insulin, refreshed every 2 days until day 10.

### Oil Red O Staining

Adipocyte differentiation was assessed by oil red O (ORO) staining. The differentiated cells were fixed in 4% paraformaldehyde for 30 minutes at room temperature and washed twice with milliQ water. Cells were then incubated in 60% isopropanol for 5 minutes and stained with freshly prepared ORO working solution for 15 minutes at room temperature, followed by three washes with milliQ water. ORO working solution was prepared by diluting 6 mL of ORO stock solution (0.5 g ORO in 100% isopropanol) with 4 mL milliQ water. Images were captured using an EXI-410 inverted microscope (ACCU-SCOPE). For quantitative analysis, dye was extracted with 100% isopropanol, and absorbance was measured using a multimode plate reader (BioTek Synergy HTX).

### Gene Expression Analysis

#### RNA isolation and PCR

Total RNA was isolated from immortalized APCs and primary ASCs using the RNeasy Mini Kit (Qiagen) according to the manufacturer’s instructions. cDNA was synthesized using the Verso cDNA Synthesis Kit (Thermo Fisher Scientific). Real-time quantitative PCR (qPCR) reactions were performed with TaqMan Fast Advanced Master Mix (Applied Biosystems). Primer/probeset IDs are in Supplemental File 1. Transcript copies were estimated from standard curves based on C_T_ values, as previously described (29).

#### Microarray analysis

Microarrays were performed by North American Genomics (Decatur, GA, USA), using the Affymetrix Clariom-S mouse HT arrays. Fifty ng of total RNA from 3 replicates of each condition were processed according to the manufacturer’s instructions for sample preparation. Data were analyzed using Affymetrix Transcription Analysis Console (TAC) software. CEL files for each experimental condition will be deposited in NCBI Gene Expression Omnibus. Gene Set Enrichment Analysis (GSEA) of Hallmark pathways was performed on normalized expression values for all detected genes from microarray experiments, as described previously (30). In addition, gene sets were created from the gene markers for each cluster described in (26). Lists of cluster markers for each gene set are in Supplemental File 2.

#### Single cell RNA sequencing analysis

The single cell RNA sequencing study and mouse model have been previously described (26) and raw files from analysis of mouse subcutaneous adipose tissue are available in the NCBI Gene Expression Omnibus GSE180880. Pseudobulk analysis was performed with Loupe software from 10x Genomics to identify differentially expressed genes in adipose stromal cells, with the following comparisons: 1) HFHS-fed (obese) versus LFLS-fed (lean) E_2_-treated mice; 2) obese E_2_ versus obese E_2_+TAM; 3) obese E_2_ versus obese EWD. The gene lists, including fold changes and p-values are found in Supplemental File 3. A Venn diagram was made using each list of genes using BioVenn (31).

To investigate Wisp2 expression in published single-cell sequencing datasets (26), quality-controlled, filtered, scaled, and SCTransformed normalized single cell RNA sequencing data were analyzed using R (v4.3.2) and the Seurat package (v5.3.0). The default assay was set to “SCT”. Wisp2 expression was visualized by cell type using Seurat’s VlnPlot function. To generate the dot plot, Wisp2 expression data were extracted using FetchData by treatment group and cell type. Mean expression and the percentage of cells expressing Wisp2 (defined as greater than 0) were calculated per treatment group and cell type. Data were plotted using the ggplot2 package (v3.5.2). Dot size is the percentage of cells expressing Wisp2 and color indicates the average expression level.

### FACS Analysis

Immortalized APCs and primary ASCs were treated with vehicle (EtOH), 10nM E_2_, 100pM 4-OH tamoxifen (TAM), 2 µg/mL recombinant mouse Wisp2 (MyBioSource), or combinations of the three. Following treatment, cells were dissociated using TrypLE™ Express (Thermo Fisher Scientific) and centrifuged at 1500 rpm for 5 minutes. Cell pellets were resuspended in FACS buffer (1x DPBS with 0.1% FBS), counted, and 1 million cells per sample were stained for 30 minutes with APC/Fire 750 anti-mouse CD24 (clone M1/69, BioLegend), PE anti-mouse Ly-6A/E (Sca-1; clone D7, BioLegend), and fixable viability dye eFluor™ 506 (Invitrogen), according to the manufacturers’ protocol. The cells were washed three times with FACS buffer and centrifuged at 1500 rpm for 5 minutes. Stained cells were analyzed and sorted using a BD FACSAria™ IIIu Cell Sorter (BD Biosciences). Flow cytometric data, including quantification of CD24⁺/Sca-1⁺ populations, were analyzed using FlowJo software (v10.10.0).

### Western Blot

Cells were lysed in RIPA buffer containing protease and phosphatase inhibitors (Roche cOmplete™ ULTRA; PhosSTOP). Protein concentrations were quantified using Pierce^TM^ BCA Protein Assay Kit (Thermo Scientific), and 50 µg of total protein per sample was loaded onto 4–15% SDS–polyacrylamide gels (Bio-Rad), followed by transfer to PVDF membranes. Membranes were blocked in 5% bovine serum albumin (BSA) in TBS-T, followed by incubation with primary antibodies (1:1000 dilution unless otherwise indicated) and HRP-conjugated secondary antibodies. Blots were developed using Pierce™ ECL Western Blotting Substrate (Thermo Fisher Scientific). Imaging was performed using the Jess™ Automated Western Blot System (Bio-Techne). Primary antibodies are listed in Supplemental File 4.

### Cell Proliferation

Cell proliferation was quantified using the Incucyte® Live-Cell Analysis System (Sartorius) within a humidified tissue culture incubator (37°C, 5% CO₂). Treatments and controls were added to experimental wells as indicated. The Incucyte® system captured images using a 10X objective in phase contrast mode in 6-12 hour intervals for 4-8 days. The Incucyte® Cell-by-Cell Analysis Software Module determined individual cells in phase channel. All experiments were performed with at least three technical replicates and repeated independently a minimum of three times.

### Statistical Analysis

Data analysis and plots were generated using GraphPad Prism 10 software, performing Šidák post-hoc comparisons, or with the R programming language version 4.5.0 using the tidyverse package. All data are expressed as mean values ± SEM with *p* values less than 0.05 considered significantly different. Two-way ANOVA, repeated measures two-way ANOVA or one-way ANOVA were performed for the analyses of differences between groups.

## RESULTS

### Estrogen and tamoxifen directly impact adipocyte precursor cells

In our previous study, we found that tamoxifen (TAM) in the presence of E_2_ depleted adipocyte progenitor cells in obese female mice compared with E_2_ alone (26), which could be due to direct or indirect actions of hormones on these cells. To investigate the direct effects of E_2_ and TAM on adipocyte precursor populations, we used three models: primary adipose stromal cells (ASCs) and immortalized adipocyte precursor cells (APCs), each isolated from subcutaneous adipose tissue of adult C57Bl/6 mice, and primary human subcutaneous ASCs. Both mouse cell lines underwent adipogenic differentiation using standard protocols reported for the widely used 3T3-L1 cell line (32), measured by oil red O accumulation (Fig 1A and Supplemental Fig 2A). The progenitor markers Cd34, Pdgfra, and Cd24a (Fig 1B-D) were elevated in proliferating and confluent cells and then decreased as differentiation progressed in both primary ASCs and immortalized APCs (Supplemental Fig 2B-C). Expression of the committed preadipocyte markers Pparg and Fabp4 (Fig 1E-F) gradually increased during the 10-day differentiation period in both cell lines (Supplemental Fig 2D-E). In primary ASCs, Esr1 gene and ERα protein levels (Fig 1G-H) increased until cells were confluent (day 0 of differentiation), decreased shortly after induction of differentiation, and then increased again by day 10. A similar gradual increase in Esr1 during differentiation also occurred in immortalized APCs (Supplemental Fig 2F).

**Figure 1.**
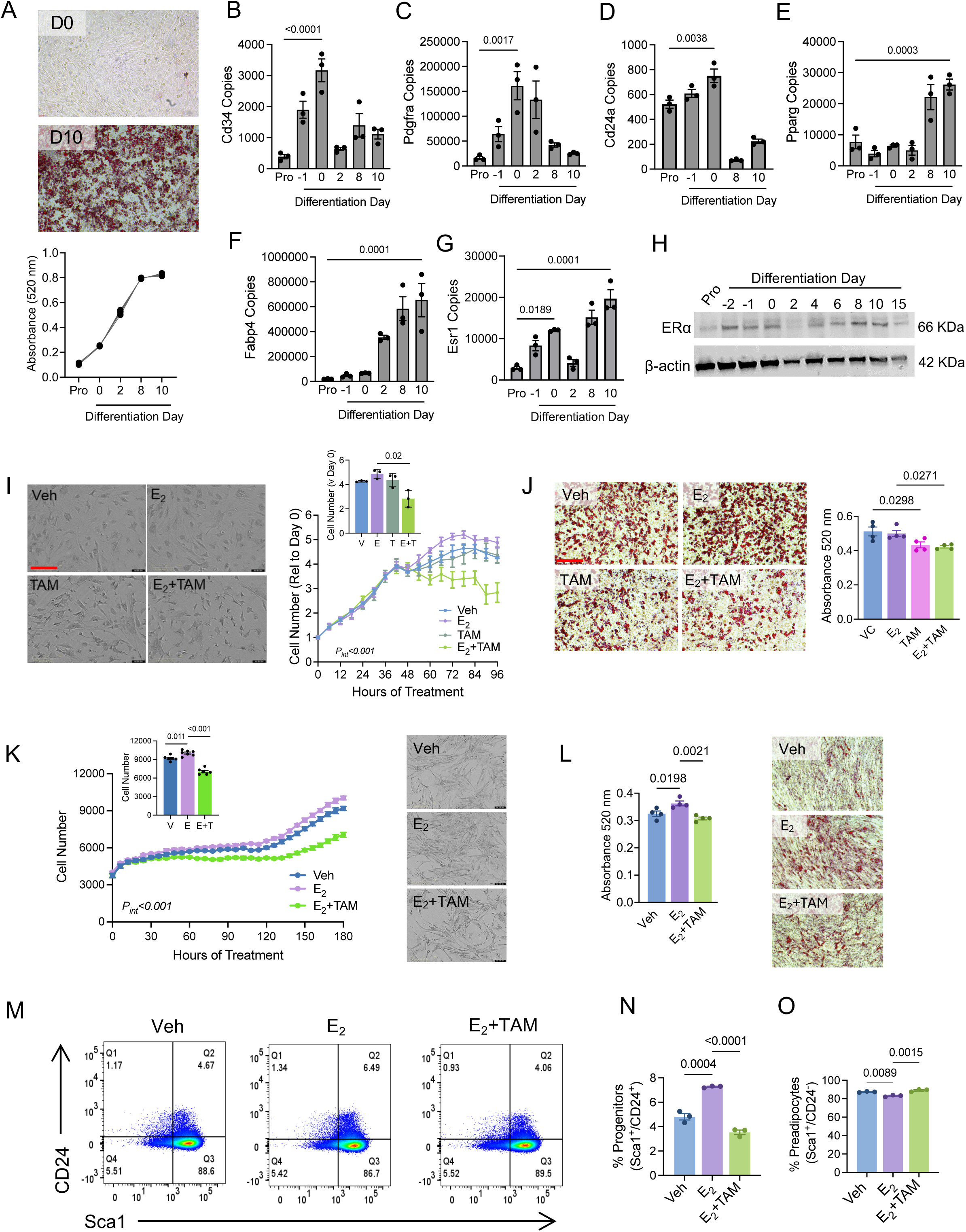
Tamoxifen inhibits effects of estradiol on subcutaneous adipocyte precursors *in vitro.* (A) Mouse subcutaneous adipose stromal cells (primary ASCs) differentiate into lipid-laden adipocytes after 10 days in culture. Images are oil red O (ORO) stained cells at day 0 (D0) or day 10 (D10) of adipocyte differentiation. Graph shows quantification of ORO accumulation over time. (B-D) Expression of adipocyte progenitor markers Cd34 (b), Pdgfra (c), and Cd24a (d) in mouse primary ASCs during proliferation (pro) or through differentiation. (E-F) Expression of preadipocyte markers Pparg (e) and Fabp4 (f) in mouse primary ASCs during proliferation (pro) or through differentiation. (G-H) Expression of Esr1 gene (g) or ERα protein (h) in mouse immortalized APCs during proliferation (pro) or through differentiation. (I) undifferentiated mouse primary ASCs treated with EtOH (Veh), E_2_, 4-OH tamoxifen (TAM) or E_2_+TAM. Images represent cells at day 4 of treatment. Scale=200µm. Inset graph represents cell number over the 4-day proliferation assay. Two-way ANOVA testing for main effects of treatment and time; interaction p<0.001. (J) ORO accumulation after 10 days of primary ASC differentiation; cells treated as in (i). Graph represents ORO absorbance on day 10. Scale=200µm. (K) undifferentiated primary human subcutaneous adipose stromal cells treated with EtOH (Veh), E_2_, 4-OH tamoxifen (TAM) or E_2_+TAM for 7.5 days. Two-way ANOVA testing for main effects of treatment or time; interaction p<0.001. Inset graph is cell number on the final day of proliferation; E_2_ versus Veh p=0.011; E_2_+TAM vs E_2_ p<0.001 by t-test. Images represent cells at day 7.5 of treatment. (L) ORO in human primary adipose stromal cells. (M) Representative FACS plots of mouse primary ASCs treated with EtOH (Veh), E_2_, or E_2_+4-OH tamoxifen (E_2_+TAM) for 48 hours and stained for Sca1 and CD24. (N-O) Percent of cells positive for Sca1 and CD24 (progenitors; n) or Sca1-positive, CD24-negative (preadipocytes, o) after treated as described in (m). T-tests determined significance.

Next, we measured expression of other ERs. In both cell lines, expression of the G-protein coupled ER, Gper1 was highest immediately before differentiation was induced at day 0 (Supplemental Fig 3A-B). Esr2 expression showed different patterns in each model, with the highest expression seen in undifferentiated immortalized APCs (Supplemental Fig 3C), but the opposite seen in primary ASCs (Supplemental Fig 3D). Of the three ERs, Esr2 had the lowest expression, detectable at ∼100 fold less than Esr1 in both cell lines (Supplemental Fig 3; Fig 1G). Human subcutaneous ASCs showed similar patterns to immortalized APCs, expressing high levels of ESR1 immediately prior to differentiation, and then again at day 9 of differentiation (Supplemental Fig 3E). GPER1 was expressed at similar levels across the differentiation time course (Supplemental Fig 3F), while ESR2 was upregulated only in fully differentiated cells (Supplemental Fig 3G).

To interrogate functional effects of hormone treatments, we measured cell proliferation and adipogenesis in response to E_2_ or TAM. In primary ASCs, E_2_ treatment slightly enhanced cell proliferation (Fig 1I). TAM alone had no effect on this outcome, but when combined with E_2_, significantly suppressed cell proliferation over a 4-day treatment period (Fig 1I). Similar effects were seen in immortalized APCs (Supplemental Fig 4A); E_2_ did not alter cell proliferation, but TAM and E_2_+TAM significantly reduced cell number over time. We then measured adipocyte differentiation using oil red O. Compared with vehicle, lipid accumulation in primary ASCs was unaffected by E_2_, but was reduced with TAM alone or with E_2_+TAM (Fig 1J). In immortalized APCs, lipid accumulation was greater with E_2_ treatment and was significantly lower with TAM or E_2_+TAM (Supplemental Fig 4B). Human ASCs responded similarly to endocrine treatments. By the end of the 7-day proliferation study, E_2_ increased cell number, and this was blocked by TAM (Fig 1K). In human ASCs, oil red O accumulation was greater after treatment with E_2,_ but this was prevented with E_2_+TAM (Fig 1L).

We and others have used Sca1 (Ly6a) and CD24 (Cd24a) to define adipocyte progenitor and committed preadipocyte populations *in vivo* (23, 26, 33). To determine how E_2_ or TAM treatments impact the proportions of these cell types *in vitro*, we analyzed Sca1^+^/CD24^+^ (progenitor) and Sca1^+^/CD24^-^(preadipocyte) populations by flow cytometry (Fig 1M; Supplemental Fig 4C). In both mouse cell lines, E_2_ treatment increased the proportion of progenitors compared with vehicle (Fig 1N; Supplemental Fig 4C). TAM inhibited the effect of E_2_, reducing the proportion of progenitors. In primary mouse ASCs, E_2_ treatment reduced the proportion of preadipocytes, while E_2_+TAM treatment increased this cell population (Fig 1O); however, hormone treatments had no impact on preadipocytes in immortalized APCs (Supplemental Fig 4C).

Estrogen is reported to facilitate proliferation of mesenchymal stem cells to support hyperplastic adipose tissue growth (34). Our previous data from obese mice implicated E_2_ in maintaining adipocyte progenitors *in vivo* during a chronic positive energy balance, as this population was depleted after giving tamoxifen to E_2_-supplemented mice (26). To determine how E_2_ and TAM influenced the renewal of adipocyte progenitor cells *in vitro*, we measured progenitors and preadipocytes in cultured primary ASCs treated with vehicle, E_2_, or E_2_+TAM after 10 days of differentiation (Fig 2A). Compared with vehicle, E_2_ treatment did not impact the proportion of progenitors (Fig 2A); however, this population was depleted in the presence of E_2_+TAM (Fig 2A). Preadipocytes were unaffected by treatment (Fig 2A).

**Figure 2.**
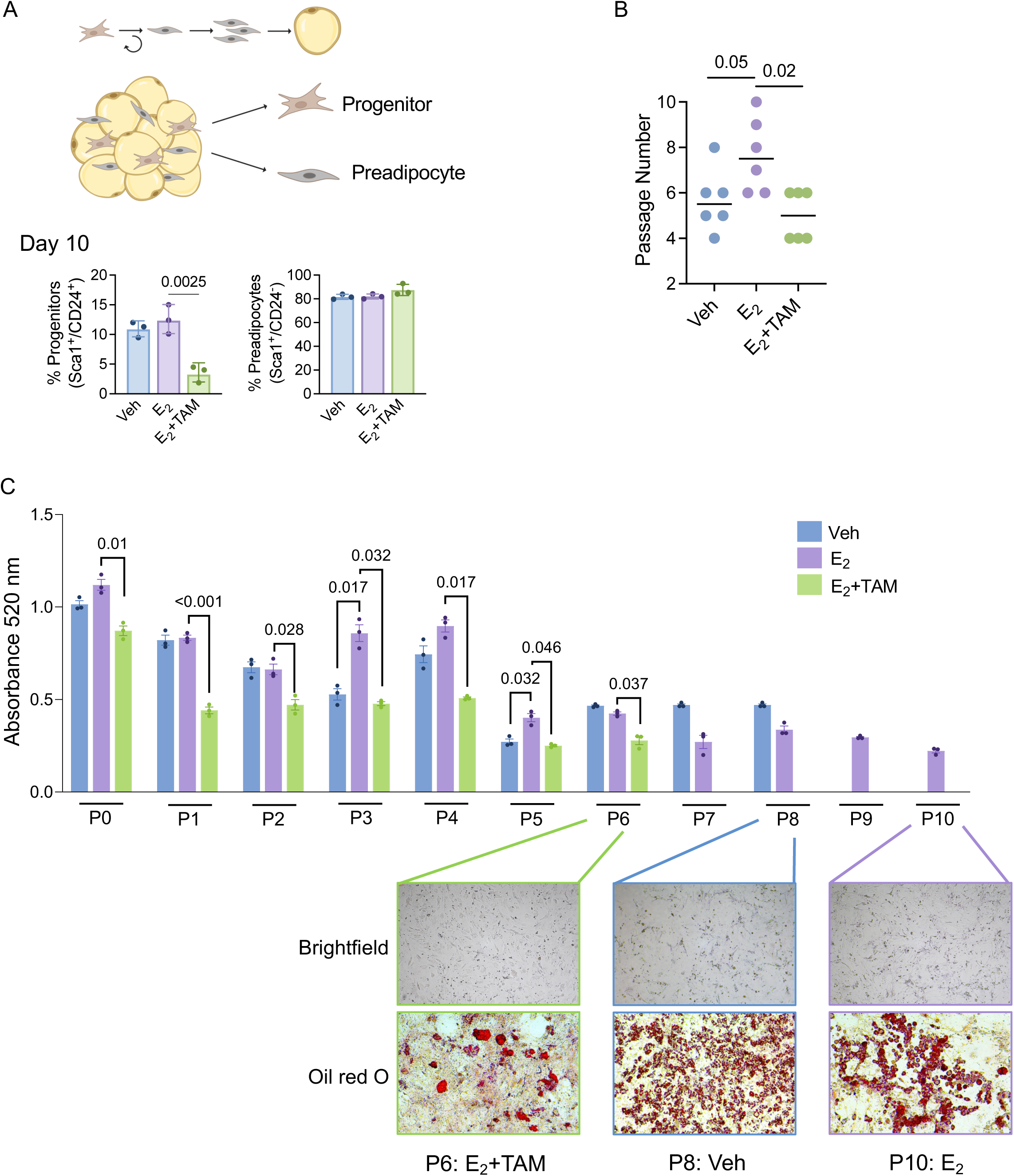
Tamoxifen reduces subcutaneous adipocyte progenitors during chronic stimulus. (A-B) Primary mouse ASCs were treated with EtOH (Veh), E_2_, or E_2_+4-OH tamoxifen (E_2_+TAM) prior to and during adipogenic differentiation for 10 days. (a). Schematic of adipogenesis was created with BioRender https://BioRender.com/az4hymd. Cells were then analyzed for Sca1+/CD24+ progenitors or Sca1+/CD24-preadipocytes by FACS (b). (C) Primary mouse subcutaneous ASCs were isolated from 6 different adult females (ages 20-24 weeks), and plated in media containing EtOH (Veh), E_2_, or E_2_+4-OH tamoxifen (E_2_+TAM). Cells were grown under these conditions and passaged each time they reached 80% confluence. The number of passages was recorded. For each treatment group, the experiment ended when cells no longer reached 80% confluence after 21 days and did not adhere to the plate with the subsequent passage, having reached the end of their proliferative lifespan. The passage number at this point was recorded for each experiment. At each passage, an aliquot of cells was plated for adipogenic differentiation and stained with oil red O (ORO). The absorbance of ORO was measured in technical triplicate wells from cells under each treatment at each passage. (D) Final passage numbers of primary ASCs were recorded under each treatment condition. Mann-Whitney test determined significance. (E) Representative quantification of ORO absorbance from key passage numbers, representing the final passage achieved for each treatment condition (E_2_+TAM=P6; Vehicle=P8; E_2_=P10). Images are phase contrast and ORO-stained plates from the respective passages.

Progenitor cells, by definition, can renew after a proliferative stimulus and can differentiate into the mature cell type for a given tissue. To evaluate each of these features, we performed a serial passaging experiment on primary mouse ASCs. After isolation from subcutaneous adipose tissue, cells were continuously cultured and passaged in hormone-depleted media that contained either E_2_, E_2_+TAM, or vehicle. Across 6 different experimental replicates, each using cells from one adult female mouse, exposure to E_2_ treatment extended the renewal of ASCs for more passages compared to vehicle treatment (Fig 2B). Addition of TAM to E_2_ prevented this renewal, and cells reached the end of their lifespan after significantly fewer passages (Fig 2B). To demonstrate that the cells remaining after chronic treatment were indeed progenitors, they were subjected to the standard adipogenesis protocol. Cells from each treatment group showed the ability to differentiate into mature adipocytes at the final passage, indicating the presence of adipocyte progenitors (Fig 2C). Together, these data demonstrate that E_2_ promotes renewal of subcutaneous adipocyte progenitors, but this protection is lost with extended TAM treatment.

### Estrogen and tamoxifen regulate Wisp2 expression in isolated adipocyte precursors

We previously identified 7 clusters of non-immune stromal cells using single cell RNA sequencing of subcutaneous adipose tissue from lean and obese female mice treated for 7 weeks with E_2_, E_2_+TAM, or estrogen deprivation/withdrawal (EWD) (26). To uncover potential mechanisms driving the *in vivo* depletion of adipocyte progenitors, we performed pseudo-bulk analysis of these clusters, comparing genes that were higher in cells from E_2_-treated obese versus lean mice to those that were lower after E_2_+TAM or EWD treatments in obese females. Wnt1-inducible signaling pathway protein 2 (Wisp2/Ccn5) met the criteria as one of two genes that were higher with E_2_ treatment (obese vs lean) and suppressed with E_2_+TAM or EWD in obese mice (Fig 3A-B). Wisp2 expression was unaffected by any treatment in lean mice. Wisp2 has been documented to maintain progenitor pools in adipose tissue of male mice (27, 28, 35). Loss of Wisp2 *in vivo* leads to adipocyte hypertrophy and glucose intolerance, similar to the phenotype that we described in female mice treated with E_2_+TAM (26). Here, we found that the population of cells previously labeled “transitional progenitors” expressed the highest levels of Wisp2 (Fig 3C), particularly in obese E_2_-treated females (Fig 3D). This cluster expresses genes common to progenitors and preadipocytes, representing a potential transitional state along the adipogenic lineage before differentiation (26). In addition to the expression level, the number of Wisp2 positive cells in the transitional progenitors, preadipocytes, and in the adipocyte regulatory (AREG) cluster, previously defined as a population that inhibits adipocyte differentiation (36), was affected by endocrine therapy. In each cluster, there were more Wisp2-positive cells in obese compared with lean E_2_-treated females, and these cells were reduced by TAM or EWD treatments (Fig 3E-G), suggesting that E_2_ and TAM might influence adipocyte precursor populations through regulation of Wisp2. In immortalized APCs, Wisp2 expression increased as cells reached confluence (day 0), then decreased after differentiation was initiated (Fig 4A-B). Similarly, in primary ASCs, Wisp2 was elevated in confluent, undifferentiated cells, decreased at day 2, and was re-expressed by day 10 (Fig 4C). In human ASCs, WISP2 was highly expressed at day 0 but was not re-expressed at day 9 of differentiation (Fig 4D). The pattern of Wisp2 expression followed that of Esr1, where gene levels peaked in confluent cells prior to induction of differentiation.

**Figure 3.**
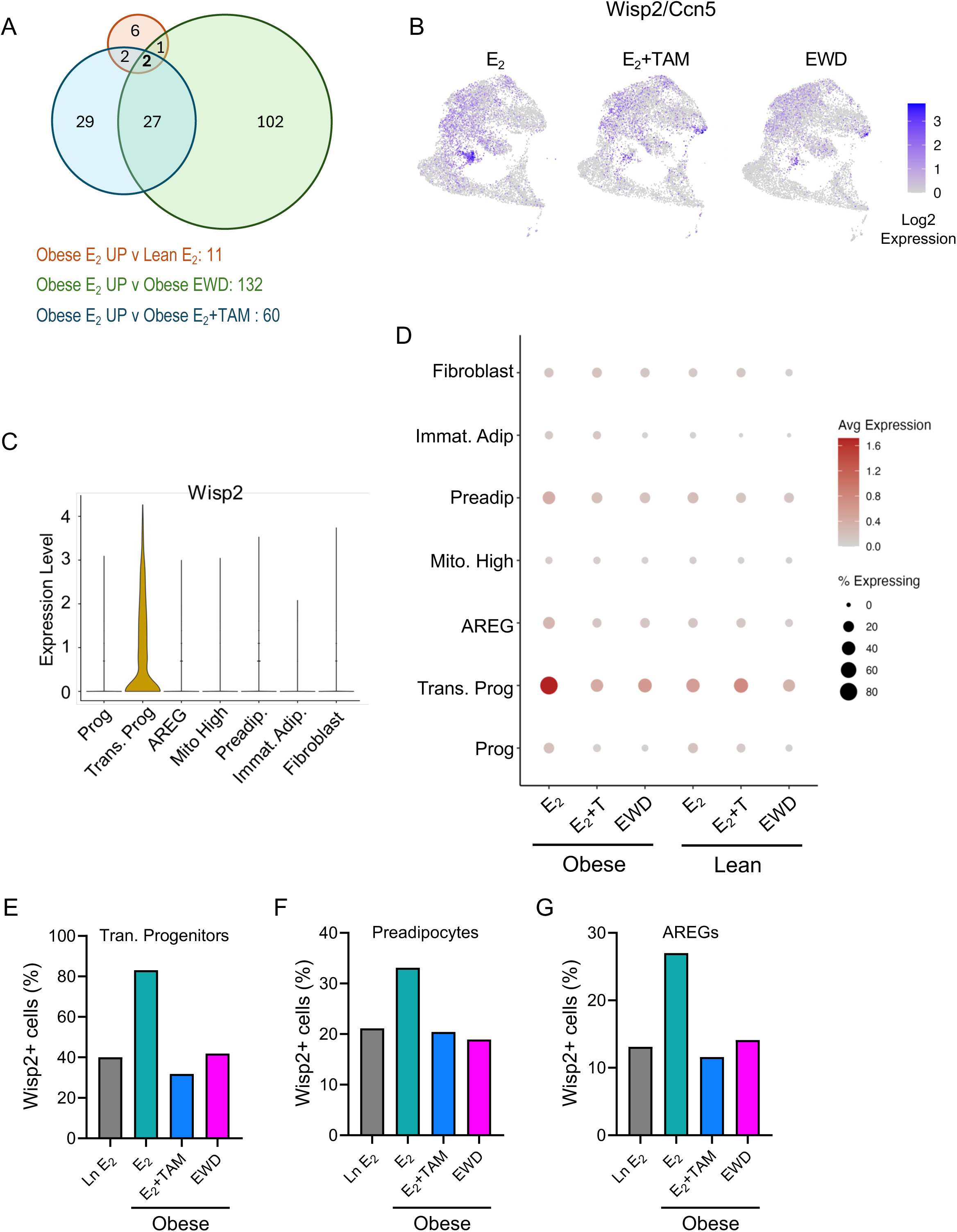
Wisp2 is downregulated by tamoxifen in mouse subcutaneous ASCs. (A) Venn diagram of genes that were higher in cells from obese versus lean female mice treated with E_2_, compared with genes that were lower after treatment of obese females with E_2_+tamoxifen (E_2_+TAM) or estrogen deprivation/withdrawal (EWD). Wisp2/Ccn5 was one of two genes in the overlap of all three gene lists. (B) UMAP showing the distribution and level of Wisp2/Ccn5 expression in primary mouse subcutaneous ASCs. (C) Violin plot showing the level of Wisp2 expression in each cluster, previously defined in Scalzo et al. (D) Bubble plot indicating the average expression level and percent positive cells for Wisp2 in each cluster and treatment group from primary mouse subcutaneous ASCs, as described in Scalzo, et al. (E-G) The percent of Wisp2 positive cells by treatment group in transitional progenitors (e), preadipocytes (f), and adipocyte regulatory cells (AREGs, g).

**Figure 4.**
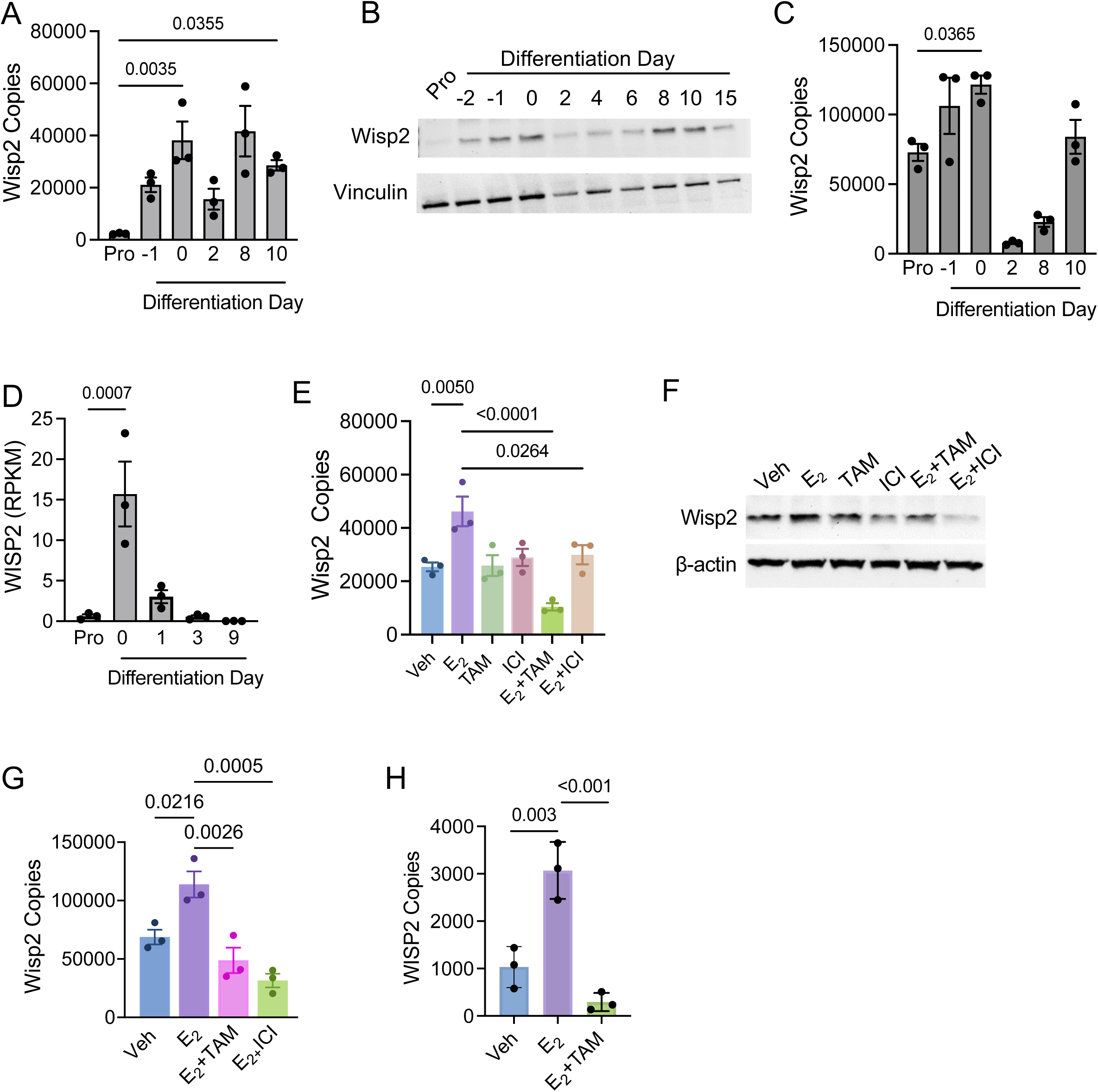
Wisp2 is regulated by E2 and tamoxifen in vitro. (A-B) Expression of Wisp2 gene (a) and protein levels (b) in mouse immortalized APCs during proliferation (pro) and differentiation. (C) Expression of Wisp2 in primary mouse ASCs during proliferation (pro) and differentiation. (D) Expression of WISP2 in primary human subcutaneous adipose stromal cells during proliferation (pro) and differentiation. (E-F) Expression of Wisp2 gene (e) and protein level (f) in immortalized mouse APCs treated with EtOH (veh), E_2_, 4-OH tamoxifen (TAM), fulvestrant (ICI), E_2_+TAM, or E_2_+ICI for 24 hours. (G) Expression of WISP2 in primary human subcutaneous adipose stromal cells treated with EtOH (Veh), E_2_, or E_2_+4-OH tamoxifen (E_2_+TAM) for 24 hours. T-tests determined significance.

Next, we investigated how estrogen signaling regulates Wisp2 *in vitro*. Treatment of immortalized APCs with E_2_ induced Wisp2 gene and protein expression (Fig 4E-F). This was prevented when ER was inhibited with TAM or the selective ER degrader, fulvestrant (ICI) (Fig 4E-F). Similar effects were seen in primary mouse and human ASCs (Fig 4G-H); E_2_ induced Wisp2, and this was inhibited by TAM. Together, these data show that Wisp2 expression profiles mirror Esr1 in cultured adipose stromal populations from mice and humans, and that Wisp2 is regulated by E_2_ and ER antagonism *in vitro* and *in vivo*.

### Wisp2 knockdown influences adipose precursor phenotypes

To determine whether Wisp2 is required for the effects of E_2_ on adipose cells *in vitro*, we reduced its expression using shRNA (Fig 5A). Compared with control cells, undifferentiated Wisp2 knockdown cells (referred to as W2KO) had lower levels of progenitor markers Cd34, Pdgfra, and Cd24a (Fig 5B-D). Overall, W2KO cells proliferated more slowly than control cells (Fig 5E; Supplemental Fig 5). In control cells, E_2_+TAM inhibited cell proliferation compared with E_2_ alone, as we saw in prior experiments. In W2KO cells, E_2_ treatment increased proliferation compared with vehicle but did not fully restore the levels seen in control cells (Fig 5E). At day 0, which is when cells have reached confluence, but differentiation has not yet been induced, the W2KO cells already displayed a rounded morphology and accumulated significantly more lipid than controls (Fig 5F), indicating precocious adipogenesis.

**Figure 5.**
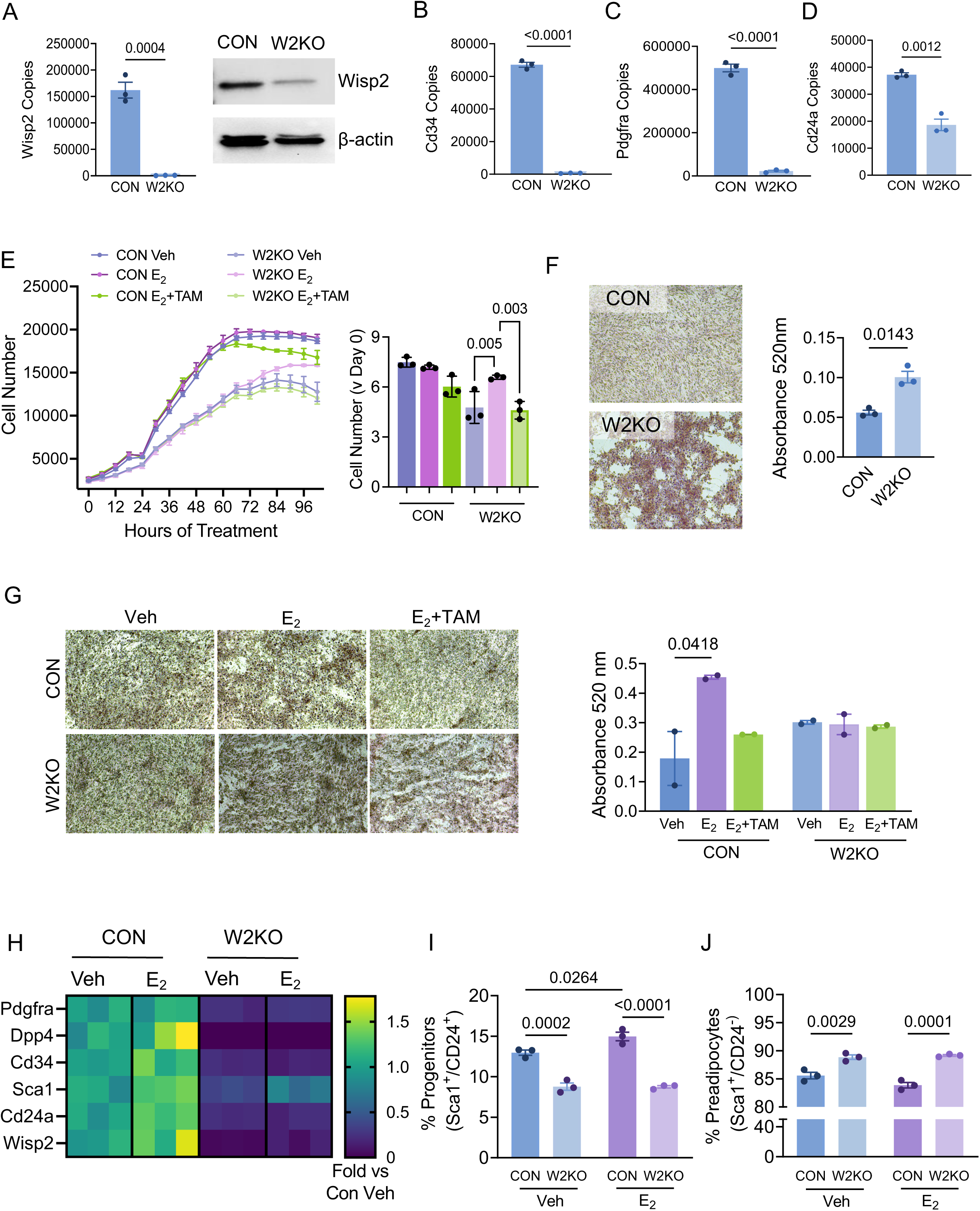
Loss of Wisp2 alters adipocyte precursor cells in vitro. (A) Expression of Wisp2 gene (left) and protein (right) in mouse immortalized APCs expressing shRNA for GFP (Con) or Wisp2 (W2KO). (B-D) Expression of adipocyte progenitor markers Cd34 (b), Pdgfra (c), or Cd24a (d) in Con and W2KO immortalized APCs during proliferation in standard growth media. (E) Cell number over 4 days in Con or W2KO populations treated with EtOH (veh), E_2_, or E_2_+4-OH tamoxifen (E_2_+TAM). 3-way ANOVA tested for main effects of time, treatment, and Wisp2 modification. (F) Oil red O (ORO) staining of Con or W2KO cells at day 0 of differentiation. Representative images on the left, quantification of ORO absorbance on the right. (G) Heatmap of expression of progenitor and preadipocyte markers in Con and W2KO cells treated with EtOH (veh) or E_2_ for 48 hours. Data are expressed as fold change versus vehicle for each gene, showing 3 replicates per group. (H-I) Quantification of percent Sca1+/CD24+ progenitors (h) and Sca1+/CD24-preadipocytes (i) in Con or W2KO cells treated with vehicle or E_2_. T-tests determined significance.

Because the W2KO cells showed enhanced differentiation even without adipogenic media, they did not remain adhered to the plates adequately to perform sustained hormone treatments. To overcome this, we repeated the differentiation assay on plates coated with type 1 collagen, a major component of adipose tissue extracellular matrix. While lipid accumulation in control cells was enhanced with E_2_ and suppressed by E_2_+TAM, there was no significant difference in lipid accumulation between treatments in W2KO cells (Fig 5G), indicating a role for Wisp2 in mediating the effects of E_2_ on adipocyte precursors.

To further investigate the effects of Wisp2 loss, we measured key markers in cells with or without E_2_. Expression of several adipocyte progenitor genes, including Pdgfra, Dpp4, Cd34, Sca1/Ly6a, Cd24a, and Wisp2 itself was higher after E_2_ treatment of control cells, but not in W2KO cells (Fig 5H; Supplemental File 5). Flow cytometry analysis revealed fewer progenitors (Fig 5I) and more preadipocytes (Fig 5J) in W2KO cells versus controls. E_2_ treatment increased progenitor proportions in control cells as expected, but this did not happen in W2KO cells (Fig 5I).

### Wisp2 treatment or overexpression alters adipogenic potential

To determine whether Wisp2 could rescue the effects of TAM on progenitors, we treated cells with exogenous Wisp2 protein and measured proliferation and differentiation. Again, as expected, E_2_+TAM treatment inhibited proliferation (Fig 6A). Wisp2 alone had no effect, but Wisp2 combined with E_2_+TAM restored proliferation (Fig 6A). Surprisingly, over the 10-day differentiation period, Wisp2 reversed the inhibitory effects of TAM and enhanced lipid accumulation in immortalized APCs and in primary ASCs (Fig 6B and Supplemental Fig 6). E_2_ treatment increased the proportion of progenitors and decreased preadipocytes (Fig 6C-E), as we consistently observed. Progenitors were significantly lower after E_2_+TAM compared with E_2_ alone, but they were maintained by treatment with E_2_+TAM+Wisp2 (Fig 6C-E).

**Figure 6.**
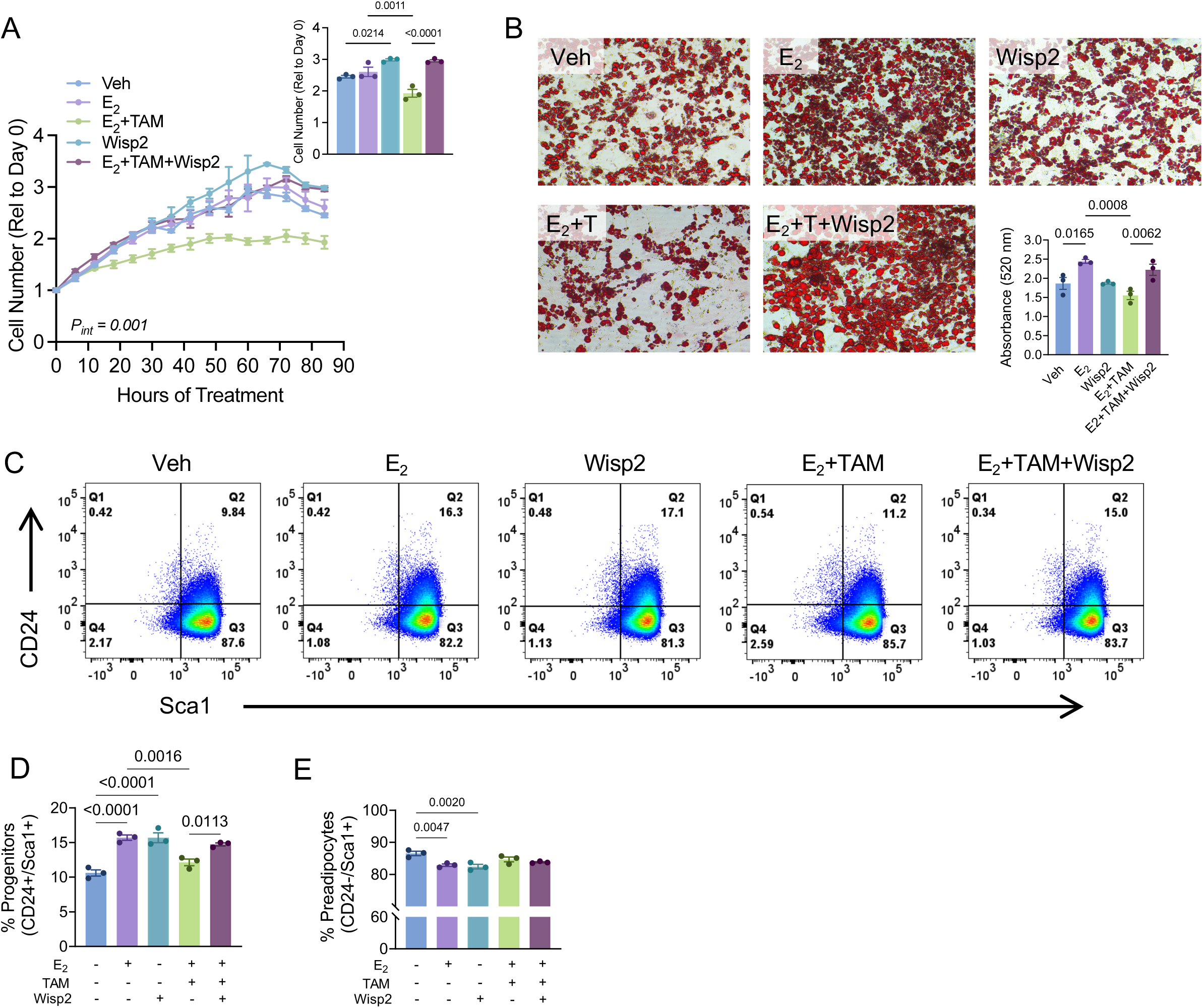
Treatment with Wisp2 reverses effects of tamoxifen on adipocyte precursors. (A) Proliferation of mouse primary ASCs during 3 days of treatment EtOH (Veh), E_2_, E_2_+4-OH tamoxifen (E_2_+TAM), exogenous Wisp2 protein (Wisp2), or E_2_+TAM+Wisp2. Two-way ANOVA tested for main effects of time and treatment; interaction p<0.001. Inset shows relative cell number at the final time point. T-test determined significance. (B) oil red O accumulation after 10 days of differentiation of cells treated as in (a). T-tests determined significance at day 10. (C-E) Representative FACS plots showing cells treated as in (a) for 48 hours and stained for Sca1 and CD24. Percent of Sca1+/CD24+ progenitors shown in (d); percent of Sca1+/CD24-preadipocytes shown in (e). T-tests determined significance. (F-K) Expression of Cd34 (f), Pdgfra (g), Sca1/Ly6a (h), Cd24a (i), Pparg (j), or Fabp4 (k) in mouse primary ASCs treated as in (a) and differentiated for 10 days. Cells were treated for the duration of the differentiation assay, replenishing media every 2 days.

To further evaluate its role as a downstream mediator of progenitor maintenance *in vitro*, we overexpressed Wisp2 in primary ASCs (Fig 7A). Consistent with prior work (37), Wisp2-overexpressing (W2OE) ASCs accumulated significantly less lipid after 10 days of differentiation and proliferated more rapidly (Fig 7C; Supplemental Fig 7) compared with controls (Fig 7B). E_2_+TAM treatment of W2OE cells slightly reduced proliferation compared with E_2_ alone, but overall, there were far more cells in any W2OE groups compared with the control groups (Fig 7C-D). W2OE increased progenitors and reduced preadipocytes as a proportion of the entire population (Fig 7E-F; Supplemental Fig 7). E_2_-treated control cells had relatively more progenitors, which were reduced with TAM, but there were no effects of E_2_ or TAM on progenitor or preadipocyte proportions in W2OE cells. Taken together, these data reinforce the role of E_2_ and TAM in regulating adipogenic cell identity and function through Wisp2.

**Figure 7.**
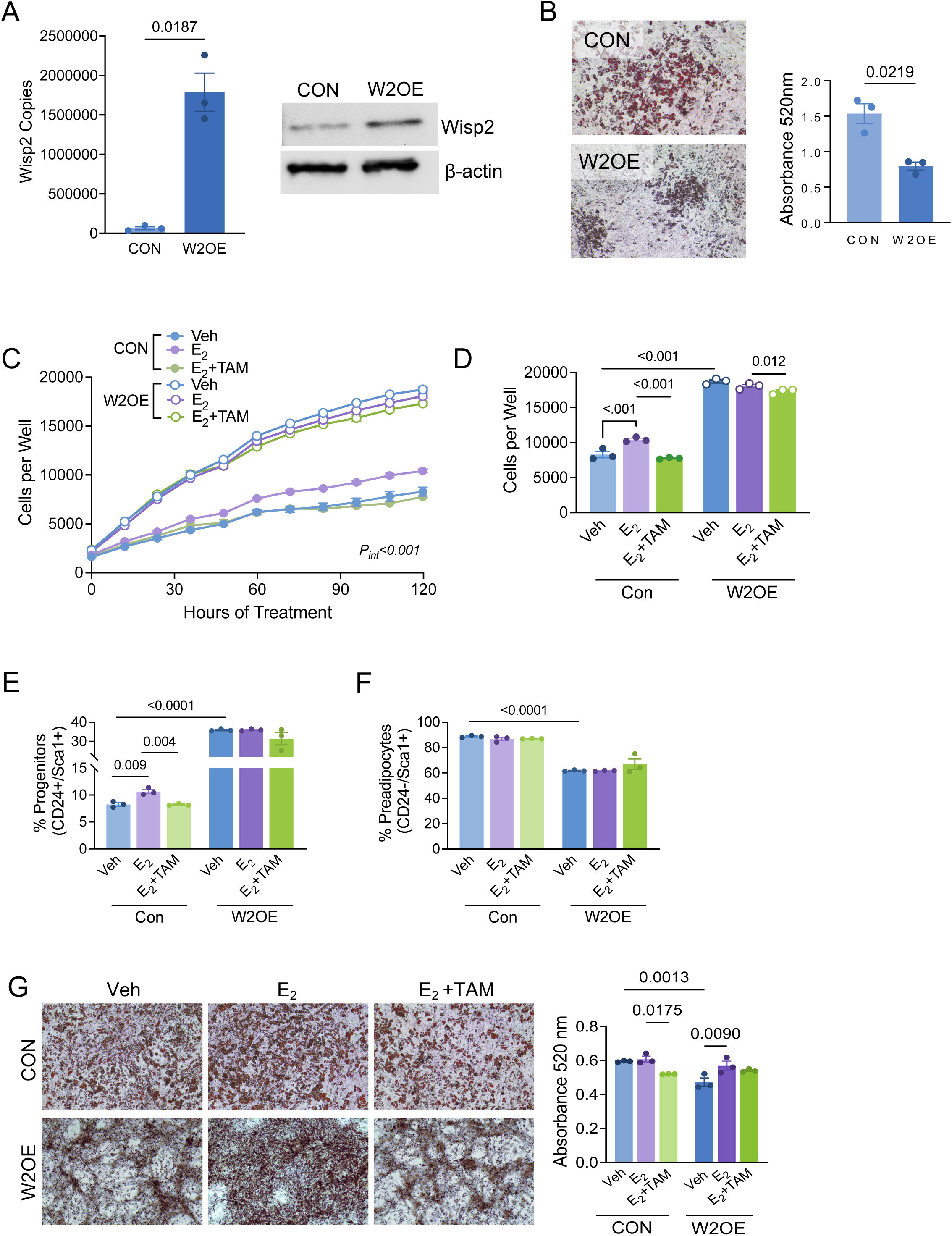
Wisp2 overexpression expands adipocyte progenitors in vitro. (A) Expression of Wisp2 gene (left) and protein (right) in primary mouse subcutaneous ASCs from wild type (Con) or Wisp2 transgenic females (W2OE) both treated with Cre adenovirus in vitro. (B) Oil red O (ORO) accumulation in control and W2OE primary ASCs differentiated for 10 days in vitro. Graph shows ORO absorbance. (C) Proliferation of control or W2OE primary ASCs over 5 days, during treatment with EtOH (Veh), E_2_, or E_2_+4-OH tamoxifen (E_2_+TAM). Three-way ANOVA tested for main effects of time, treatment, or cell genotype. Interaction p-value <0.001. (D) Quantification of cell number after 5 days of proliferation as shown in (c). Statistical analysis is two-way ANOVA testing for main effects of treatment or cell genotype, with post-hoc analyses. (E-F) Quantification of Sca1+/CD24+ progenitors (e) or Sca1+/CD24-preadipocytes (f) after 2 days of treatment with EtOH (Veh), E_2_, or E_2_+TAM measured by FACS. (G) ORO accumulation in CON or W2OE cells treated with EtOH (Veh), E_2_, or E_2_+TAM for 10 days during differentiation. Graph represents quantification of ORO absorbance at day 10. T-tests determined significance.

### Endocrine therapies and Wisp2 alter adipocyte precursor trajectories

To evaluate and compare changes in gene expression profiles, we performed transcriptomic and pathway analyses following treatment of cells with E_2_ or TAM, or in cells with or without Wisp2. Compared with control cells, E_2_ treatment had minor effects, but induced genes related to IFNα or IFNγ signaling, while inhibiting expression of mitotic spindle genes and those related to UV response (Supplemental File 6). Treatment with E_2_+TAM increased genes related to IFNα and IFNγ, similar to E_2_, but effects on other pathways were distinct. E_2_+TAM treatment increased TNFα and TGFβ signaling and inflammatory response genes, while reducing expression of genes associated with epithelial to mesenchymal transition (EMT). Knockdown of Wisp2 had similar effects to E_2_+TAM, as expected, increasing genes related to TNFα and TGFβ signaling, and reducing expression of EMT-associated genes, compared with control cells. Despite those similarities, IFNα and IFNγ signaling genes were altered in the opposite direction as with E_2_+TAM. These data suggest that some effects of TAM are mediated through Wisp2, but in other cases, TAM might have ER agonist activity in adipocyte precursors.

Interestingly, exogenous Wisp2 treatment and Wisp2 overexpression affected few of the same pathways (Supplemental File 6). Wisp2 overexpression (W2OE) increased genes associated with IFNγ and TNFα signaling, much like treatment with E_2_, but also increased EMT and inflammatory response pathways. Exogenous Wisp2 treatment suppressed TNFα- and p53 pathway genes, and increased adipogenesis-related genes, aligned with its augmentation of lipid accumulation. Treatment of cells with exogenous Wisp2 would presumably only activate cell surface Wnt signaling pathways and would not function in the cytosol to prevent Pparg induction (35). When added to E_2_+TAM, exogenous Wisp2 almost completely reversed the effects of TAM on gene expression, suppressing IFNα, IFNγ, and TNFα signaling, hypoxia, and oxidative phosphorylation networks, and increasing EMT and mitotic spindle genes.

Although GSEA is informative, many genes are found in more than one Hallmark pathway and no single pathway captured the expression profiles of the distinct adipose stromal cell populations. To further explore these subtle but important cell states, we created gene sets (markers) using differentially expressed genes in each distinct cluster within the stromal fraction of subcutaneous adipose tissue of female mice in our prior study (26) and analyzed each cell population and treatment together using GSEA (Fig 8A; Supplemental File 7). E_2_ treatment modestly increased transitional progenitor markers. Addition of TAM to E_2_ interfered with this phenotype, suppressing progenitor and preadipocyte markers. E_2_+TAM treatment also upregulated genes associated with the inhibitory AREG cluster (36). Knockdown of Wisp2 (W2KO) phenocopied E_2_+TAM treatment, reducing progenitor, transitional progenitor, and preadipocyte markers. W2KO cells also had elevated expression of immature adipocyte markers, consistent with the precocious lipid accumulation that we observed. Cells overexpressing Wisp2 (W2OE) had the opposite expression profile as W2KO, with lower immature adipocyte markers and higher levels of transitional progenitor, AREG, progenitor, and preadipocyte genes. Exogenous Wisp2 treatment alone (Ex-W2) had more subtle effects, suppressing immature adipocyte and progenitor markers, but increasing genes associated with transitional progenitors. Addition of exogenous Wisp2 to E_2_+TAM reversed effects of TAM, lowering genes associated with immature adipocytes and increasing markers of progenitors, transitional progenitors, and preadipocytes. Altogether, these data demonstrate a role for E_2_ and TAM in regulating adipose stromal cell populations, dependent in part upon Wisp2 (Fig 8B).

**Figure 8.**
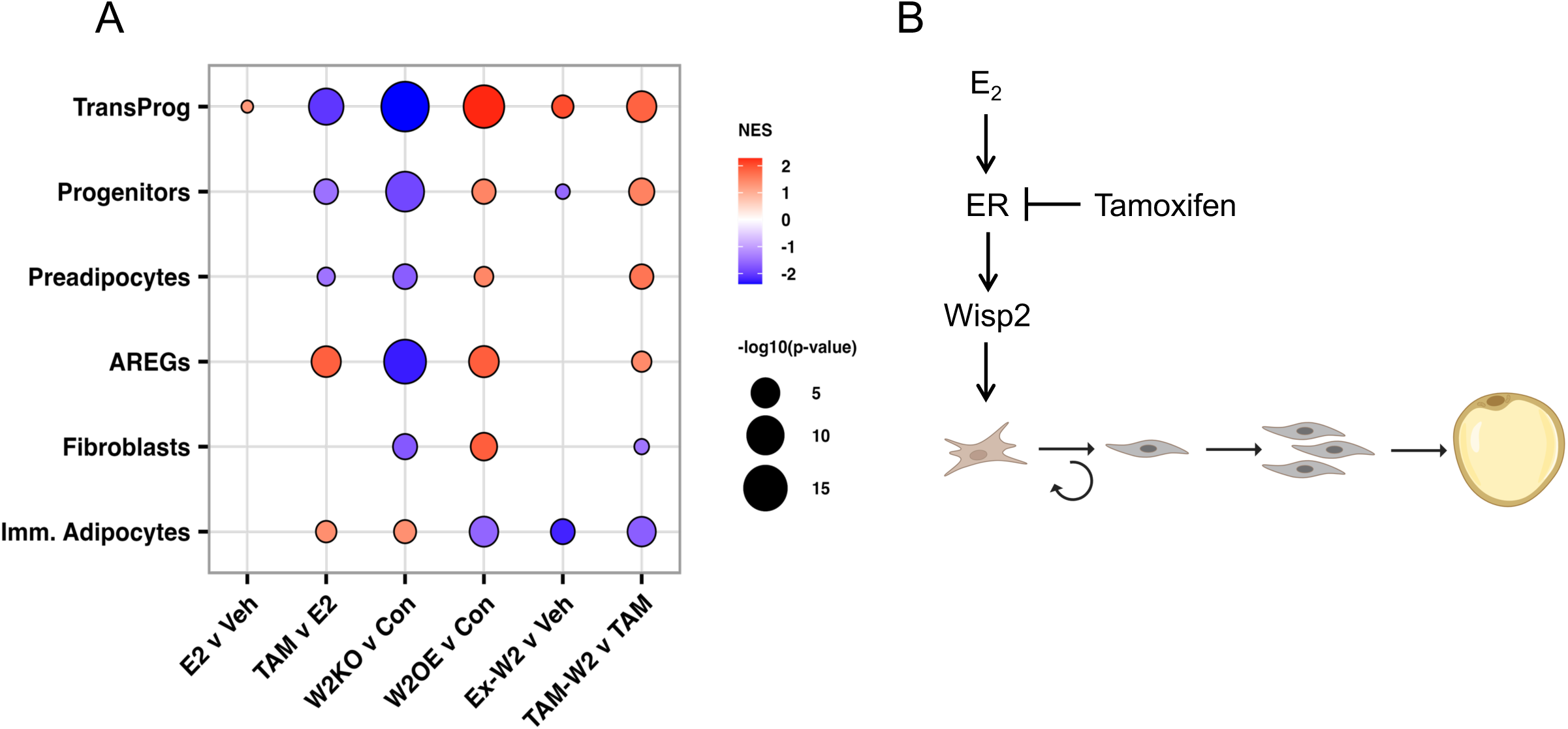
Modulation of estrogen signaling or Wisp2 alters adipose stromal cell populations. (A) Bubble plot of gene set enrichment analysis performed on transcriptomics data collected from the following groups: Lanes 1, 2, 3, 5, 6 represent immortalized mouse APCs treated with EtOH (Veh) versus E_2_; E_2_+4-OH tamoxifen (TAM) versus E_2_; expressing Wisp2 shRNA (W2KO) versus GFP shRNA (Con); treated with exogenous Wisp2 versus PBS+BSA (Veh); or treated with E_2_+TAM+Wisp2 versus E_2_+TAM. Lane 4 represents primary mouse ASCs expressing Wisp2 (W2OE) versus Cre-treated controls (Con). Normalized enrichment scores of each gene set, along with the log10 p-values are depicted by the bubbles. Gene sets were derived from prior annotation of clusters present in subcutaneous adipose stromal cells isolated as CD45-/CD31-/CD29+ and used for single cell RNA sequencing as in Scalzo et al. (B) Working model representing the inhibitory effect of TAM on E_2_-induced Wisp2 expression. Wisp2 mediates the maintenance of adipocyte progenitors downstream of E_2_ and ER activation, which is disrupted with TAM.

## DISCUSSION

Menopause or estrogen-modulating therapies alter metabolism and adipose tissue distribution (6–8, 38–40). A lower capacity for subcutaneous adipose expansion can occur after menopause in women, and associates with an excess risk for cardiometabolic diseases (41). Clinical data and our published preclinical study show prevalent metabolic dysfunction following breast cancer endocrine therapy, including ectopic fat deposition and insulin resistance (3, 26). We previously found that TAM treatment in obese mice depleted the Sca1^+^/CD24^+^ progenitors in subcutaneous adipose tissue (26); cells that are highly adipogenic and responsible for renewal of preadipocyte pools (23). This observation reveals adipose tissue as a primary target of endocrine therapy, as ERα is expressed in both undifferentiated precursors and mature adipocytes (42). In this study, we use “precursors” to define the heterogeneous population of cells that contains both progenitors and committed preadipocytes; each of which can be evaluated by flow cytometry based on cell surface marker expression. Estrogen signaling promotes the proliferation and maintenance of progenitors *in vivo*, particularly within obesogenic environments (24, 34). Because the loss of adipocyte progenitors that we previously observed *in vivo* could result from direct or indirect effects of endocrine therapy on adipose cell types, the objective of this study was to determine how E_2_ and TAM directly impact the diverse adipocyte precursor population, and to define underlying mechanisms that potentially mediate the effects.

Our data show that both E_2_ and TAM act directly on adipocyte precursors to influence their gene expression profiles and function. We show that E_2_ supports the enrichment of progenitors *in vitro* after adipogenic differentiation and enables extended renewal and proliferation of primary adipose stromal cells over time; effects that are inhibited by TAM. These results are consistent with the depletion of adipocyte progenitors observed in mice lacking Esr1 in this cell population (42). In AdipoTrak ERα knockout mice, in which Esr1 was selectively knocked down in progenitors, the overall number of this cell type was reduced in subcutaneous depots from males and females (42). Further investigation revealed that stromal cells lacking Esr1 shifted their fate, adopting a vascular smooth muscle cell phenotype with elevated expression of TGF² signaling intermediates. The effects of TAM that we identified here are consistent with prior work showing that loss of ERα signaling in adipose tissue drives reprogramming of white adipocyte progenitors, altering their fate and reducing adipose tissue plasticity (43, 44). One prior study evaluated the effects of tamoxifen on breast adipose stromal cells *in vitro*, showing that TAM reduced proliferation, increased apoptosis, and partially prevented the induction of Pparg and Lpl genes during adipogenic differentiation (45). Our results are consistent with those findings, however we used the active metabolite 4-OH tamoxifen in combination with E_2_ and administered tamoxifen at lower concentrations than in the published study (45).

Here, we identified Wisp2 as a mediator of TAM’s effects on adipocyte precursors, measuring lower Wisp2 expression in a subcutaneous adipose progenitor population from TAM-treated compared with E_2_-treated obese female mice. In cultured adipocyte precursors from mice and humans, E_2_ treatment increased Wisp2 expression, and this was prevented with TAM, reinforcing a direct role for ER in Wisp2 gene regulation. While E_2_ can induce Wisp2 in breast cancer cells (46, 47), this has not been shown in adipose tissue stromal cells and reveals a potentially widespread mechanism of estrogen-dependent Wisp2 regulation. Wisp2 acts through at least two mechanisms to maintain adipocyte precursor cells in their undifferentiated state. As a secreted adipokine, it stimulates canonical Wnt/β-catenin signaling to directly repress Pparg transcription and block the expression of adipogenic genes (48). Additionally, in the cytosol Wisp2 binds to and forms a complex with the zinc-finger transcription factor Zfp423, preventing its nuclear entry and subsequent Pparg activation, thereby restricting adipocyte commitment and differentiation (37). Here, we found that reduction of Wisp2 levels *in vitro* attenuated cell proliferation, depleted progenitor pools, and accelerated adipogenic differentiation. These findings are consistent with work by Hammarstedt et al (35), which demonstrated that Wisp2 knockdown in 3T3L-1 mouse preadipocytes induced spontaneous adipogenesis in the absence of differentiation stimuli, characterized by C/ebpο, Pparγ and C/ebpα activation. In our study, E_2_ treatment consistently increased progenitor proportions in immortalized and primary adipose precursors, but it was unable to expand progenitor cells in Wisp2-deficient APCs, suggesting that Wisp2 mediates the maintenance of progenitor pools by E_2_.

Interestingly, the consequences of Wisp2 overexpression and exogenous Wisp2 treatment were not identical. Both exogenous Wisp2 treatment and Wisp2 overexpression prevented the inhibitory effects of TAM on cell proliferation, suggesting that the pro-proliferative effect of Wisp2 on adipocyte precursor cells is primarily mediated by cell surface Wnt pathway activation. While exogenous Wisp2 treatment restored lipid accumulation in TAM-treated cells, Wisp2 overexpression impaired lipid accumulation. The divergent effects of Wisp2 on adipogenesis in our study are consistent with its distinct mechanisms of action at the cell surface and in the cytosol, both of which help to maintain adipocyte progenitor pools. Together these observations reveal Wisp2 as an essential regulator of adipocyte progenitor cell fate and renewal rather than simply a cell type-specific marker.

Transcriptional analysis showed similarities between tamoxifen-treated and W2KO populations, revealing that both perturbations shifted gene profiles away from transitional progenitor and preadipocyte markers, and towards immature adipocyte profiles. Wisp2 overexpression and exogeneous Wisp2 treatment both increased the expression of characteristic transitional progenitor and preadipocyte genes compared to their respective controls, suggesting the maintenance and proliferation of undifferentiated ASCs and APCs. Administration of exogeneous Wisp2 to TAM-treated APCs reversed the changes in genes related to pro-inflammation, cellular stress, hypoxia, and oxidative phosphorylation. Studies have shown that these pathways are disrupted in ERα knockout adipocytes or APCs, which have features of metabolic and inflammatory dysfunction (43, 49). Together, these studies demonstrate a role for adipocyte ERα in adipose tissue inflammation and metabolism, and our work adds to these observations by demonstrating changes in the stromal cells. The suppression of stress and inflammatory pathway genes that we observed also indicates that TAM might elicit the development of inflammatory and dysfunctional adipocyte progenitors; an effect that remains to proven *in vivo*.

In this study, we focused on the actions of Wisp2 *in vitro* to determine how it directly impacts adipocyte precursor cells downstream of estrogen signaling. It is not yet known if Wisp2 overexpression would overcome the effects of TAM *in vivo*. To this end, we bred transgenic mice expressing a conditional Wisp2 construct to the widely used Pdgfra-Cre model, with the goal of inducing Wisp2 in adipocyte precursors during TAM treatment *in vivo*. However, mice expressing both Pdgfra-Cre and conditional Wisp2 (i.e. adipocyte precursor Wisp2-overexpressing mice) were unable to survive any surgical procedures due to a severe bleeding defect, characterized by the inability of blood to clot. It is reported that up to 20% of adult hematopoietic cells derive from Pdgfra-positive precursors (50), suggesting that Wisp2 overexpression in such cells might alter their ability to appropriately differentiate into crucial cell types that contribute to wound healing (51). Regardless, this unexpected phenotype limited our ability to model endocrine therapy *in vivo* because we were unable to perform ovariectomy surgeries; a key feature of our estrogen manipulation model (26).

In summary, our data show that TAM interferes with the ability of E_2_ to protect adipocyte progenitor populations in subcutaneous adipose tissue, partly mediated by suppression of Wisp2. Taken together with our prior study demonstrating the loss of adipocyte progenitors after TAM treatment of obese mice, these data suggest a direct mechanism of E_2_ and TAM action on precursor populations from subcutaneous adipose tissue. Future studies will define whether interventions that overcome these inhibitory effects can restore healthy adipose tissue expansion during endocrine therapy to attenuate the development of metabolic disease in females.

## Acknowledgements

This work was supported by the Stephenson Cancer Center (SCC), the Harold Hamm Diabetes Center (HHDC), and University of Oklahoma Health in Oklahoma City, Oklahoma, the HHDC/SCC Postdoctoral Fellowship (NST), as well as the Health and Environmental Sciences Institute THRIVE grant (EAW).

## FIGURE LEGENDS

**Supplemental File 1:** Primer/probe catalog numbers used for qPCR.

**Supplemental File 2:** Lists of cluster markers used to create adipose stromal GSEA gene sets.

**Supplemental File 3:** Differentially expressed genes from pseudobulk analysis of adipose stromal cells in lean and obese mice.

**Supplemental File 4:** Primary antibodies used for analyses.

**Supplemental File 5:** Raw data values for heatmap in Figure 5H.

**Supplemental File 6:** GSEA analysis of Hallmark Pathways.

**Supplemental File 7:** GSEA analysis of adipose stromal clusters.

