## Supplemental Figures for "Tamoxifen Targets Wisp2 to Impair Subcutaneous Adipocyte Progenitor Self-Renewal and Adipogenic Differentiation"

Supplemental Fig 1

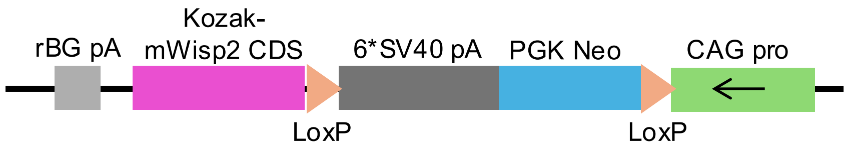


**Supplemental Figure 1.** Transgene construct used to generate conditional Wisp2-overexpressing adipose stromal cells. Wisp2 transgenic mice were created by The Taconic-Cyagen Model Generation Alliance.

Supplemental Figure 2

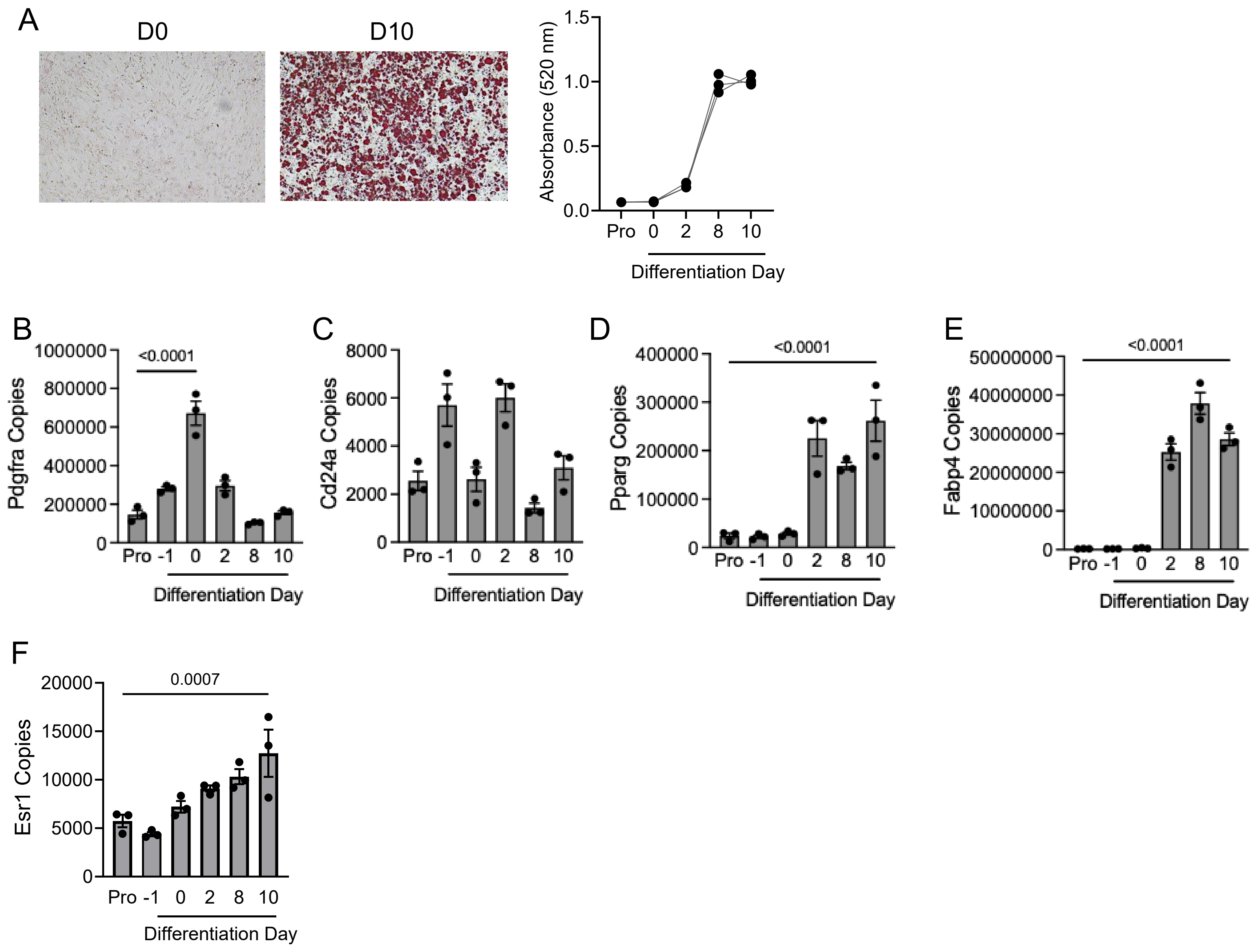


**Supplemental Figure 2. Characterization of immortalized mouse subcutaneous adipocyte precursor cells.** (A) Mouse immortalized adipocyte precursors (immortalized APCs) differentiate into lipid-laden adipocytes after 10 days in culture. Images are oil red O (ORO) stained cells at day 0 (D0) or day 10 (D10) of adipocyte differentiation. Graph shows quantification of ORO accumulation over time. (B-C) Expression of adipocyte progenitor markers Pdgfra (b), and Cd24a (c) in immortalized APCs during proliferation (pro) or through differentiation. (D-E) Expression of preadipocyte markers Pparg (d) and Fabp4 (e) in immortalized APCs during proliferation (pro) or through differentiation. (F) Expression of Esr1 gene in immortalized APCs during proliferation (pro) or through differentiation. For b-f, one way ANOVA determined significance, followed by post-hoc comparisons.

Supplemental Figure 3

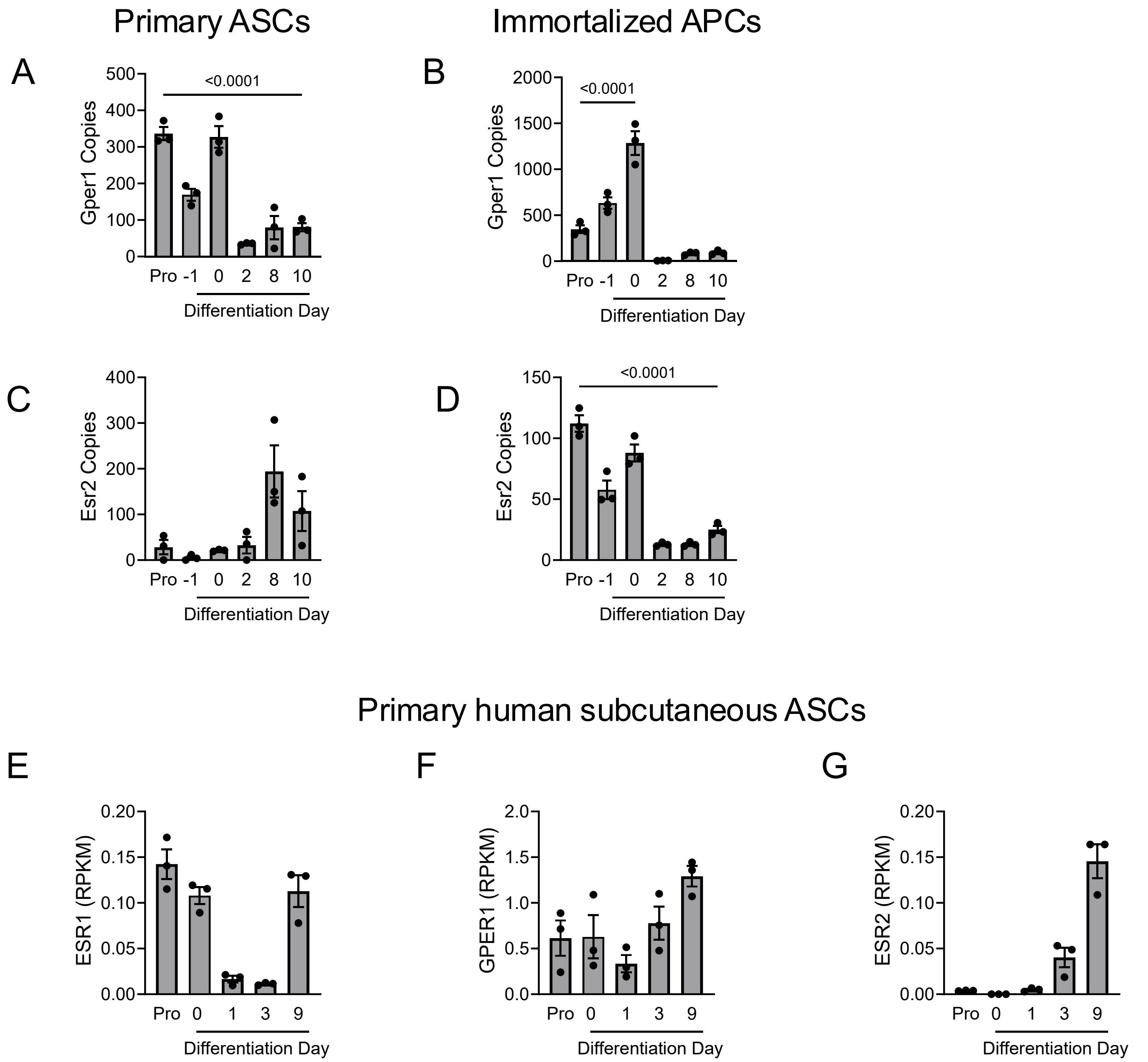


**Supplemental Figure 3. Characterization of estrogen receptors in cultured cells.** (A-B) Expression of Gper1 in primary ASCs (a) or immortalized APCs (b). (C-D) Expression of Esr2 in primary ASCs (c) or immortalized APCs (d). One way ANOVA followed by post-hoc comparisons determined significance. (E-G) Expression of ESR1 (e), GPER1 (f), or ESR2 (g) in primary human subcutaneous adipocyte precursor cells during proliferation (pro) or differentiation.

Supplemental Figure 4

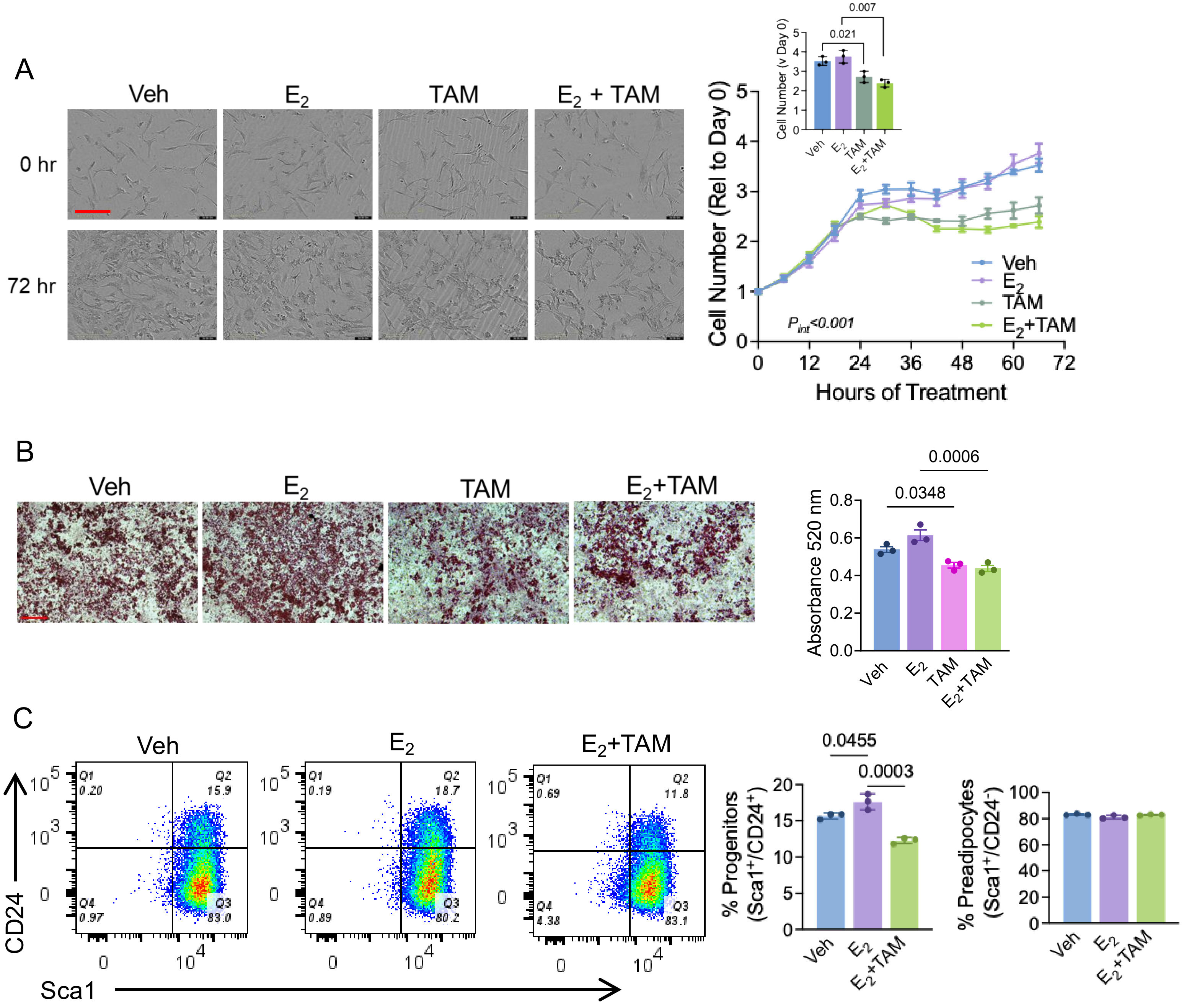


**Supplemental Figure 4. Estradiol and tamoxifen influence proliferation and differentiation of mouse immortalized APCs.** (A) undifferentiated mouse immortalized APCs treated with EtOH (Veh), $E_2$, 4-OH tamoxifen (TAM) or $E_2$+TAM. Images represent cells at day 0 and day 3 of treatment. Graph represents cell number over the 3-day proliferation assay. Scale=400µm. Two-way ANOVA testing for main effects of treatment and time; interaction $p<0.001$. (B) Oil red O (ORO) accumulation after 10 days of immortalized APC differentiation; cells treated as in (a). Graph represents ORO absorbance on day 10. T-tests determined significance. Scale=200µm. (C) Representative FACS plots of mouse immortalized APCs treated with EtOH (Veh), $E_2$, or $E_2$+4-OH tamoxifen ($E_2$+TAM) for 48 hours and stained for Sca1 and CD24. (D-E) Percent of cells positive for Sca1 and CD24 (progenitors; n) or Sca1-positive, CD24-negative (preadipocytes, o) after treated as described in (m). T-tests determined significance.

Supplemental Figure 5

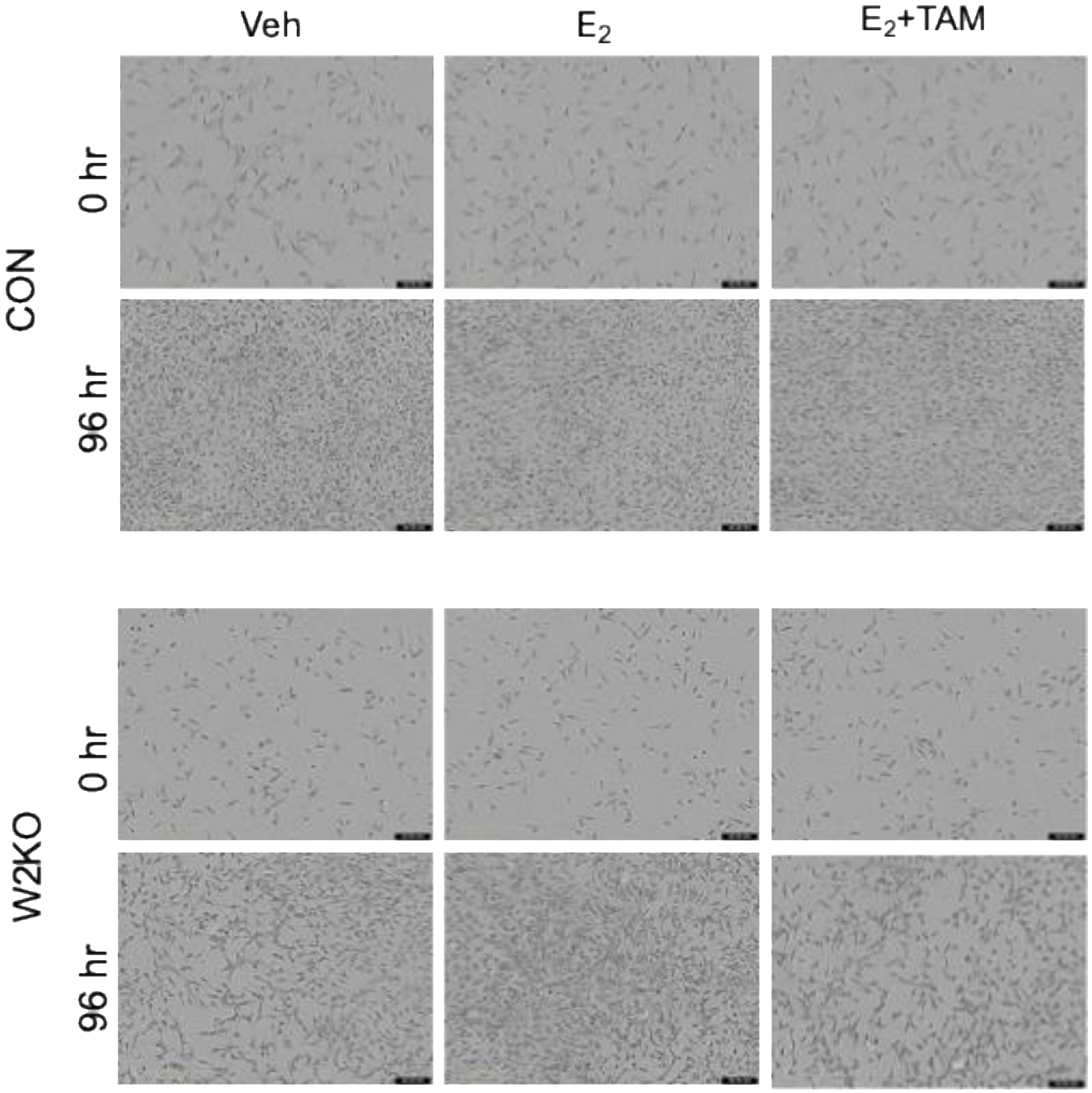


**Supplemental Figure 5. Loss of Wisp2 influences cell proliferation.** Representative images of control or Wisp2 knockdown (W2KO) cells at 0 hours or 96 hours of treatment with EtOH (Veh), $E_2$, or $E_2$+4-OH tamoxifen ($E_2$+TAM). Data are presented in Figure 5.

Supplemental Figure 6

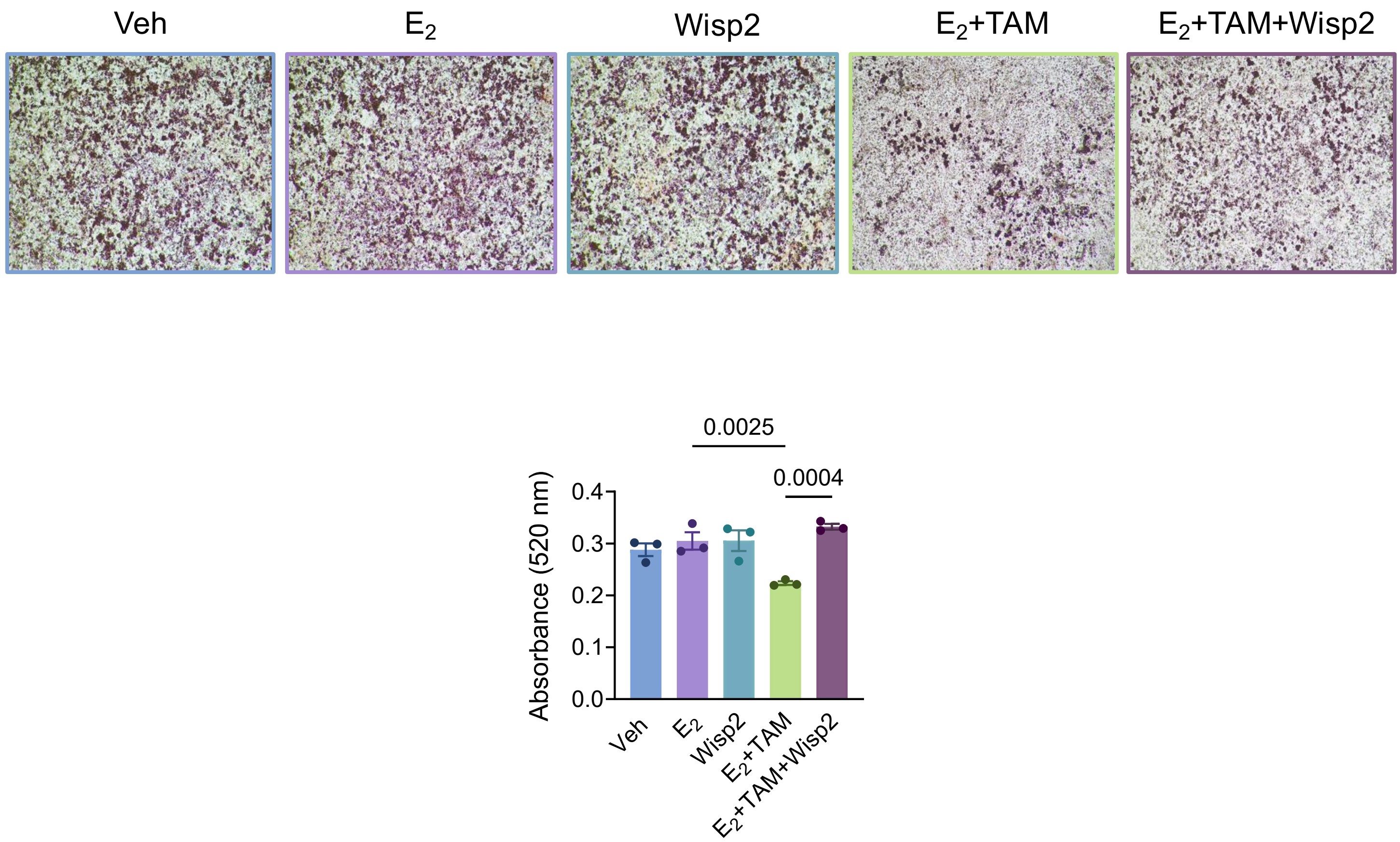


**Supplemental Figure 6. Wisp2 restores lipid accumulation in tamoxifen-treated immortalized APCs.** Representative images of oil red O (ORO) accumulation are shown for each treatment. Quantification of ORO at day 10 is shown in the graph. T-tests determined significance.

Supplemental Figure 7

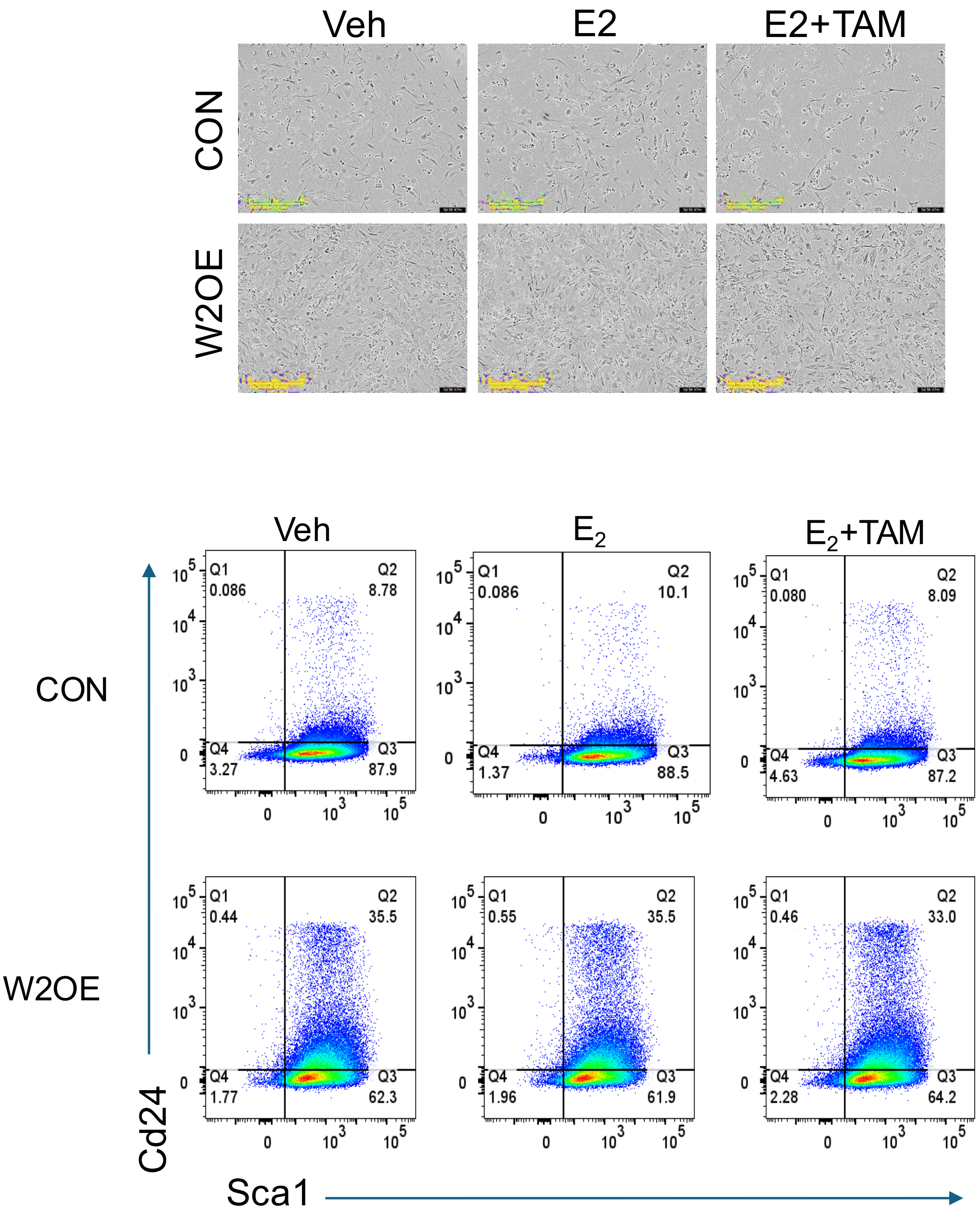


**Supplemental Figure 7. Cell Proliferation and FACS images of Control and Wisp2-overexpressing cells.** (A) Images of CON and W2OE cells treated with EtOH (Veh), $E_2$, or $E_2$+4OH-tamoxifen ($E_2$+TAM) after 5 days of proliferation assay, quantified in Figure 7C-D. (B) Representative FACS plots of cells described in Figure 7E-F.
