## Supplemental File 1 for "Tamoxifen Targets Wisp2 to Impair Subcutaneous Adipocyte Progenitor Self-Renewal and Adipogenic Differentiation"

**Supplemental File 1**: TaqMan primers used for qPCR experiment

| **Vendor** | **Catalog Number** | **Primer** | **Assay ID** | **Species** | **Method** |
| --- | --- | --- | --- | --- | --- |
| Fisher Scientific | 4453320 | Pdgfra | Mm00440701_m1 | Mouse | TaqMan |
| Fisher Scientific | 4331182 | PPARγ | Mm01184322_m1 | Mouse | TaqMan |
| Fisher Scientific | 4331182 | Wisp2 | Mm00497471_m1 | Mouse | TaqMan |
| Fisher Scientific | 4331182 | Fabp4 | Mm00445878_m1 | Mouse | TaqMan |
| Fisher Scientific | 4331182 | Esr2 | Mm00599821_m1 | Mouse | TaqMan |
| Fisher Scientific | 4331182 | Esr1 | Mm00433149_m1 | Mouse | TaqMan |
| Fisher Scientific | 4448892 | Dpp4 | Mm01329189_m1 | Mouse | TaqMan |
| Fisher Scientific | 4453320 | CD34 | Mm00519283_m1 | Mouse | TaqMan |
| Fisher Scientific | 4453320 | CD24a | Mm00782538_sH | Mouse | TaqMan |
| Fisher Scientific | 4448892 | WISP2 | Hs01031984_m1 | Human | TaqMan |
